# Distinct airway progenitor cells drive epithelial heterogeneity in the developing human lung

**DOI:** 10.1101/2022.06.13.495813

**Authors:** Ansley S. Conchola, Tristan Frum, Zhiwei Xiao, Peggy P. Hsu, Renee F.C. Hein, Alyssa Miller, Yu-Hwai Tsai, Angeline Wu, Kamika Kaur, Emily M. Holloway, Abhinav Anand, Preetish K. L. Murthy, Ian Glass, Purushothama R. Tata, Jason R. Spence

## Abstract

Recent advances using single cell genomic approaches have identified new epithelial cell types and uncovered cellular heterogeneity in the murine and human lung (1). Here, using scRNA-seq and microscopy we identify and describe a secretory-like cell that is enriched in the small airways of the developing human lung and identified by the unique co-expression of *SCGB3A2/SFTPB/CFTR*. To place these cells in the hierarchy of airway development, we apply a single cell barcode-based lineage tracing method track the fate of *SCGB3A2/SFTPB/CFTR* cells during airway organoid differentiation *in vitro* (2). Lineage tracing revealed that these cells have distinct developmental potential from basal cells, giving rise predominantly to pulmonary neuroendocrine cells (PNECs) and a subset of multiciliated cells distinguished by high *C6* and low *MUC16* expression. We conclude that *SCGB3A2/SFTPB/CFTR* cells act as a progenitor cell contributing to the cellular diversity and heterogeneity in the developing human airway.

**SIGNIFICANCE STATEMENT:** The current study identifies a novel secretory cell type that is present predominantly in the small airway of the developing human lung. These secretory cells are defined by co-expression of *SCGB3A2/SFTPB/CFTR*, and functional studies show that this cell gives rise to pulmonary neuroendocrine cells and a sub-population of multiciliated cells, thereby leading to cellular heterogeneity.

## INTRODUCTION

For decades, scientists have relied on morphology, histologic approaches, and low throughput methods to visualize a limited number of proteins or mRNA transcripts to identify and characterize human tissues. However, our understanding of the highly specialized cells within human organs has been improved by recent technological advances in single cell genomic and high-resolution spatial microscopy and transcriptomic approaches to interrogate tissue composition and organization (1, 3–23). Only in the past few years, for example, troves of single cell and spatial transcriptomic data within the lung field have described the vast cellular heterogeneity in the adult human lung at homeostasis (3, 10, 16, 24–26), or in various pathologic states, including (but not limited to) asthma (26), pulmonary fibrosis(15, 17, 20), cystic fibrosis (6), and infections such as SARS-CoV-2 (8, 27). Several studies have also revealed the complexities of the developing human lung (1, 13, 14, 21, 23, 28). This new information has shed light on cellular heterogeneity within the human lung, how cells change fate throughout the process of development, and has identified interesting new cell types or states that are uniquely human with no obvious correlate in the well-studied murine lung (10, 24). However, studying human organs and tissue has inherent limitations based on ethical considerations and limited access, often making functional follow-up studies difficult, particularly in the case of human development.

Our group recently published a single cell RNA sequencing (scRNA-seq) dataset characterizing the human fetal lung epithelium from 8 – 21 weeks post-conception (1) and used this dataset to benchmark an *in vitro* lung organoid model (29) that differentiates bud tip progenitor cells to airway cell types, including basal stem cells. During our analysis of the human fetal lung scRNA-seq data, we identified a cohort of cells expressing a unique combination of genes encoding secreted proteins including *SCGB3A2* and *SFTPB*. Recently, this unique expression profile has been identified in cells in the small airways and terminal respiratory bronchioles of adult human lungs (10, 19, 24, 30); however, human fetal *SCGB3A2/SFTPB* cells also express *CFTR*, which was not described in these adult cells. Furthermore, *SCGB3A2/SFTPB/CFTR* cells in fetal lung appear to lack genes canonically associated with secretory (i.e., club, goblet), alveolar (i.e., alveolar epithelial type II cells), or ionocyte cell types. Based on their specificity to lung development, anatomic location, and gene expression signature, we refer to this population as <u>F</u>etal <u>A</u>irway <u>S</u>ecretory (FAS) cells.

Given the unreported nature of these cells, it was previously unclear where they emerge in the airway hierarchy. The goal of the current manuscript is to characterize the FAS cell population within the human fetal lung and determine the functional role of this population during lung development. We predicted it might serve as a progenitor during development given its unique presence in the fetal lung. Here, *in situ* analyses suggest FAS cells arise during branching morphogenesis and are located to small cartilaginous and non-cartilaginous (middle airways) of the fetal lung. Lineage tracing of *in vitro* lung organoid models and functional experiments using organoids suggest FAS cells are a distinct airway progenitor from the basal cell lineage, giving rise to pulmonary neuroendocrine cells (PNECs) and a subset of multiciliated cells defined by *C6* expression, while basal cells are an airway progenitor for club cells and multiciliated cells defined by *MUC16*. Additionally, these multiciliated cell subsets have regional specificity within the lung airway, in correlation with basal and FAS cells. Collectively, this study reveals complex cellular and functional heterogeneity within the fetal lung epithelium during development.

## RESULTS

### scRNA-seq identifies a novel epithelial cell during development defined by co-expression of *SCGB3A2/SFTPB/CFTR*

Original scRNA-seq data from our group (1) composed of 8 to 21 post conception weeks (PCW) human lung tissue dissections of trachea, proximal airway, and distal airway, and was reanalyzed for epithelium only (Figure 1A). The data was analyzed using Seurat (31–34) and epithelial populations were identified using Louvain clustering (35) followed by visualization via UMAP (36, 37) (Figure 1A). Cell populations were annotated using cohorts of genes associated with different cell types in the human fetal lung as previously described (1) (Figure S1A). This analysis identified many of the previously characterized populations of lung epithelial cell types including bud tip progenitors, the cell types that give rise to all epithelial cells of the lung, bud tip adjacent cells, basal cells, the resident stem cell of the lung, secretory club and goblet cells which provide secretions that lubricate and line the airways, multiciliated cells which help clear debris and move secretions, and pulmonary neuroendocrine cells (PNECs) which have sensory capabilities and secrete peptides into the environment (Figure 1A, S1A). We also identified a robust population of cells that we call ‘Fetal Airway Secretory’ (FAS) cells, that was positioned on the UMAP embedding between the ‘bud tip adjacent’ cluster, ‘basal cell’ cluster, and ‘club-like secretory’ cluster (Figure 1A, pink cluster). This cluster has a unique gene expression profile that included co-expression of *SCGB3A2, SFTPB*, and *CFTR* (Figure 1B). The co-expression of these three markers together is unique and has not been reported before. Of interest, while SCGB3A2 expression has been associated with club cells, SFTPB with alveolar type II cells, and CFTR with ionocytes and other cells in the adult airway (6, 16, 38), FAS cells did not express other club (*SCGB1A1, SCGB3A1*), alveolar (*SFTPC, SFTPA1*), or ionocyte (*FOXI1*) markers (Figure S1B, Table S1). We also noted that *SCGB3A2, SFTPB*, and *CFTR* were not uniformly expressed across the cluster, so we computationally extracted and re-clustered these cells to interrogate heterogeneity within this population (Figure 1C). We observed 5 subclusters, with subcluster 2 possessing the most specific co-expression of *SCGB3A2, SFTPB*, and *CFTR* (Figure 1C). Other canonical secretory genes from both proximal and distal cell types were analyzed for expression which were minimally expressed in subcluster 2 (Figure S1C). Notably, *RNASE1* was also enriched in subcluster 2 which has recently been reported in an *SCGB3A2*^*+*^/*SFTPB*^+^/*SCGB1A1*^-^ cell population in the adult (10). The FAS cell sub-cluster (Cluster 2 in Figure 1C-D) was analyzed by sample age and lung region to determine the proportion of FAS cells from each (Figure 1E-F). Cells that were derived from small airways contributed 73% of cells in subcluster 2, 19% of cells were from tracheal samples, and 8% from distal lung samples. By age, 57% of cells come from 8-15 PCW samples; the 18 PCW sample contributes 39% and 19-21 PCW samples contribute 3.5% to the cluster. A similar analysis was performed on the entire FAS cell cluster prior to sub-clustering (i.e. from Figure 1A), and supported the trends observed by analyzing sub-clustered FAS cells (Figure S1D-E).

**Figure 1:**
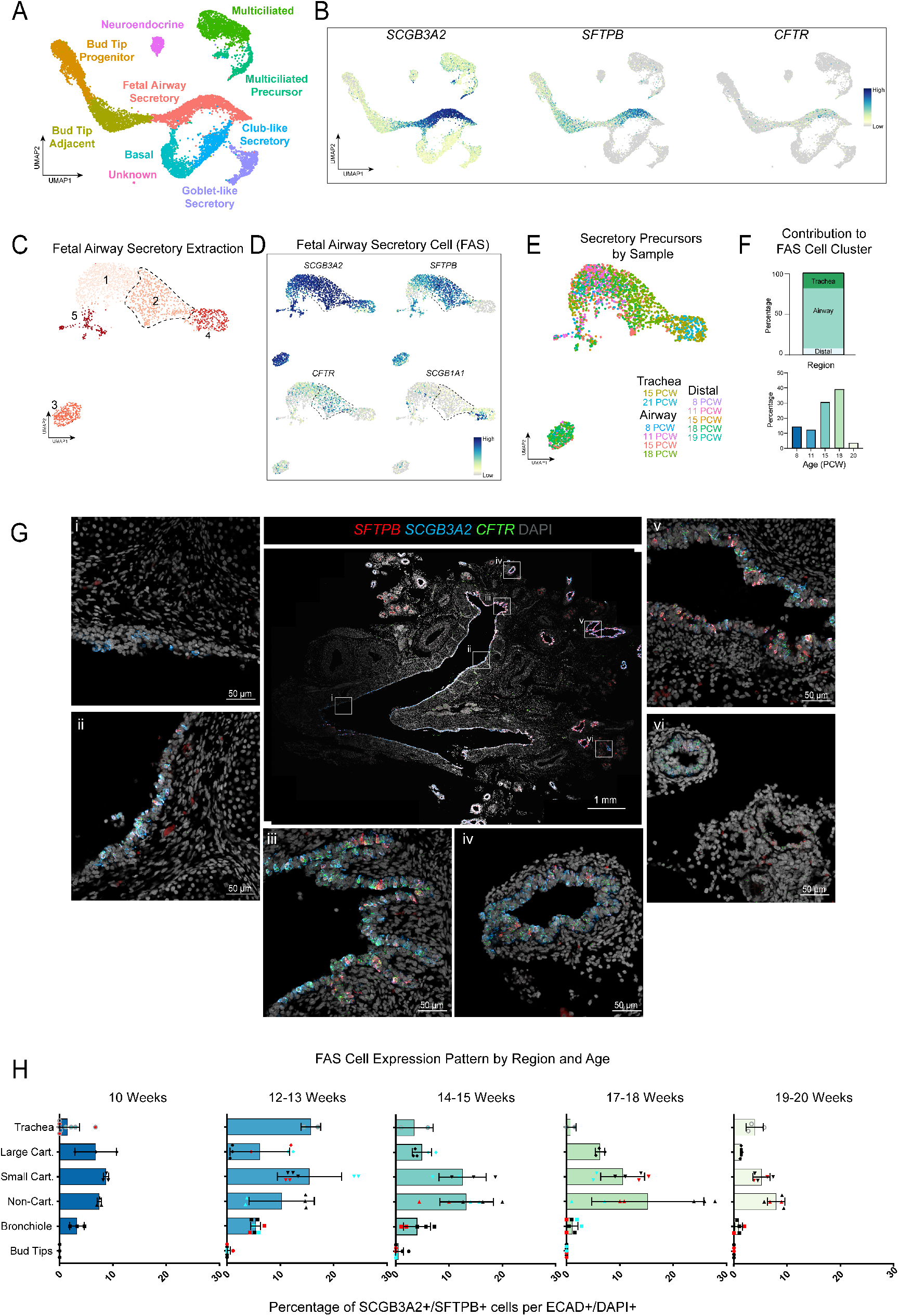
scRNA-sequencing identifies *SCGB3A2*^*HI*^*/SFTPB*^*HI*^*/CFTR*^*HI*^ secretory precursor in middle airways. (A) UMAP cluster plot of scRNA-seq data from human fetal lung epithelium. 6 biologically distinct samples were sequenced at PCW ages 8, 11, 15, 18, 19, 21 from distinct regions of trachea, airway, or distal lung. Each dot represents a single cell and cells were computationally clustered based on transcriptional similarities. Clusters are colored and labeled by cell type, which were determined based on expression of canonical cell-type markers displayed in the dot plots in Fig. S1. (B) UMAP feature plots of highly expressing genes *SCGB3A2, SFTPB*, and *CFTR*, which co-express predominantly in the Fetal Airway Secretory Cluster from Fig. 1A. (C) UMAP cluster plot of Fetal Airway Secretory Cluster from Fig. 1A computationally extracted and re-clustered. Each dot represents a single cell and cells were computationally clustered based on transcriptional similarities. The plot is colored and numbered by cluster. Cluster 2 is outlined to correspond with feature plots in Fig. 1D. (D) Feature plots of *SCGB3A2, SFTPB, CFTR*, and *SCGB1A1. SCGB3A2, SFTPB*, and *CFTR* co-express predominantly in subcluster 2 (outlined). *SCGB1A1* is only highly expressed in subcluster 4. (E) UMAP cluster plot of subclustered FAS cells by sample. Each dot represents a single cell and cells were computationally clustered based on transcriptional similarities. 6 biologically distinct samples were sequenced at PCW ages 8, 11, 15, 18, 19, 21 and distinct regions of trachea, airway or distal lung were sequenced among the samples. (F) Quantification of samples’ contributions to FAS cell subcluster 2 are reported as percentages by lung region (top panel) or sample age (bottom panel). (G) Representative fluorescence *in situ* hybridization (FISH) images of 11.5 PCW lung for FAS cell markers *SCGB3A2, SFTPB*, and *CFTR*. Boxed regions within middle panel correspond to regions shown from trachea (i), bronchus (ii, iii), small cartilaginous (v), non-cartilaginous (iv) and distal bud tip (vi). (H) Quantification of SCGB3A2^+^/SFTPB^+^ expressing cells among 100 ECAD^+^/DAPI^+^ cells imaged across distinct lung regions (trachea, large cartilaginous, small cartilaginous, non-cartilaginous, bronchioles, bud tips) and ages (10 – 20 PCW).

To validate the presence of FAS cells in tissue, we performed fluorescence *in situ* hybridization (FISH) for all three genes on human fetal lung samples ranging from 8 to 20 PCW. Abundant *SCGB3A2*^*+*^*/SFTPB*^*+*^*/CFTR*^*+*^ co-expressing cells were identified at all stages examined, and representative low and high-magnification images from12 PCW lung tissue sections from trachea, bronchi, large and small cartilaginous and distal bud tips are shown (Figure 1G). Co-expressing cells appear to cluster together in the airways, decreasing in abundance distally throughout the epithelium. To support scRNA-seq and FISH data, we also identified FAS cells using SCGB3A2/SFTPB co-immunofluorescence (co-IF) across stages and regions and quantified the spatiotemporal location of FAS cells within the epithelium (Figure 1H). We observed changes in co-staining both in cell number and spatial distribution across time that were consistent with our scRNA-seq analysis. In 10 PCW samples, 27% of epithelial cells co-expressed SCGB3A2 and SFTPB. FAS cells were most abundant (comprising 53% of epithelial cells) in 12-13 PCW samples, with the number of cells decreasing over time with 19% of epithelial cells co-expressing SCGB3A2/SFTPB at 20 PCW (38% in 13-14 PCW, 33.8% in 17-18 PCW). The regional distribution of cells demonstrated that SCGB3A2^+^/SFTPB^+^ cells are most abundant in the small cartilaginous (52.4%) and large non-cartilaginous airways (54.2%) with far fewer cells present in more proximal (25.3% trachea, 25.5% large cartilaginous) or distal (14% bronchiole, 0% bud tip) regions (Figure 1H). Together, this analysis suggests FAS cells are enriched in the middle airways of the developing lung, and decrease in abundance by 20 PCW.

### An *in vitro* bud tip organoid differentiation model to functionally interrogate FAS cells

To generate FAS cells for functional studies *in vitro*, we used primary human bud tip organoids (BTOs) (1, 29, 39) which can be grown as a homogenous population of SOX2^+^/SOX9^+^ BTPs (1, 29) or can be differentiated into airway using a 21-day “dual SMAD activation/inhibition” (DSA/DSI) Airway Differentiation Paradigm, which includes 3 days of DSA followed by 18 days of DSI (Figure S2A, see methods, and (1, 40)). This Airway Differentiation Paradigm has previously been shown to be robust in inducing proximal airway cell types including basal, multiciliated, neuroendocrine, and secretory cells, and prior scRNA-seq suggested that induced airway organoids possessed abundant FAS cells (1, 40).

However, the timing when FAS cells emerge and their perdurance during airway differentiation was not interrogated further. To better understand when FAS cells are present during airway differentiation, we performed a time course using qRT-PCR on BTOs throughout the DSA/DSI protocol (Figure S2B). BTOs were collected at 30 min, 3hr, 24hr, and 3 days during DSA, and 30 min, 3hr, 24hr, 11 days, and 17 days during DSI. Consistent with our previous reports, *TP63*, a marker of basal cells and early airway differentiation, increased during the first 24hr and remained elevated (1), whereas FAS cell markers (*SCGB3A2, SFTPB*) decreased during DSA, but increased after switching to DSI media and maintained high expression throughout the remainder of the time series (Figure S2B). To further interrogate the presence of FAS cells in airway organoids, we carried out FISH with co-IF on 21-day airway organoids and confirmed the presence of *SCGB3A2*^+^/SFTPB^+^/*CFTR*^+^ FAS cells (Figure S2C). This led us to infer that FAS cells emerge early in the differentiation paradigm and were maintained throughout its duration, thereby allowing us to utilize this model to interrogate FAS cell function.

### Single cell barcoded lineage tracing suggests that FAS cells are progenitors for PNEC and multiciliated cells

Next, we leveraged airway differentiation of the BTO model to determine if FAS cells have the potential to give rise to other airway epithelial cells during airway differentiation. We implemented a barcode-based lineage tracing technique, called CellTagging (2) which utilizes a complex lentiviral library of unique and heritable barcodes affixed to the 3’ UTR of GFP enabling us to tag and track clones of individual cells using scRNA-seq. BTOs were differentiated into Airway Organoids using the DSA/DSI Airway Differentiation Paradigm to generate an initial population of FAS cells and basal cells at 21 days (Figure 2A). At the 21-day timepoint, organoids were needle-passaged to shear them into small fragments, subsequently transduced with the lentiviral CellTagging V1 Library (2), and replated in 3D matrix to reform organoids. GFP expression could be detected within 24hr and was robustly expressed in reformed organoids on day 7 (Figure S2D). qRT-PCR of infected organoids and uninfected controls demonstrated no differences in expression of many canonical lung epithelial markers (Figure S2E) suggesting viral transduction had no effect on airway differentiation potential. 7 days after infection and passaging, infected cultures were dissociated into single cell suspension and subject to fluorescence activated cell sorting (FACS) to isolate GFP^+^ cells, demonstrating that approximately 15% of cells were GFP^+^ (Figure S2F, average of 3 experiments). CPM is a well-established cell surface marker of bud tip progenitor (BTP) cells (40, 41), so we negatively sorted CPM^HI^ cells to remove any undifferentiated BTPs from the culture (Figure S2F) which could further differentiate and confound analysis. Additionally, only 2% of GFP^+^/CPM^-^ cells were replated in 3D matrix with a limited dilution to reduce the probability that multiple cells from the same clone are present in the starting culture (see ‘Methods’). Thus, majority of organoids in this culture should derive from a single uniquely tagged FAS or basal cell (Figure 2A). After replating sorted cells, organoids reformed and were allowed to expand for 30 days at which point they were collected for scRNA-seq (Figure 2A). Of 8,256 total cells sequenced, 43.4% of cells contained CellTags (n = 3,583) with similar distribution throughout all clusters (30-50%) (Figure S2G). Cell types were identified using a gene module-based cell scoring method which evaluates *in vitro* cells for expression of a panel of genes generated from the fetal epithelial scRNA-seq data (Figure 2B, S1A, S2H-I) (see ‘Methods’). The cell types identified in this organoid model shared a high degree of transcriptional similarity to *in vivo* basal cells, secretory cells, multiciliated precursors (deuterosomes), multiciliated cells and PNECs (Figure 2B-C, Table S2). One cluster remained unidentified by the cell type scoring; given this cluster expressed early markers of PNEC differentiation (*ASCL1*^*HIGH*^) (42–44) and its proximity to the PNEC cluster, we assigned this cluster a PNEC precursor identity (pink cluster).

**Figure 2:**
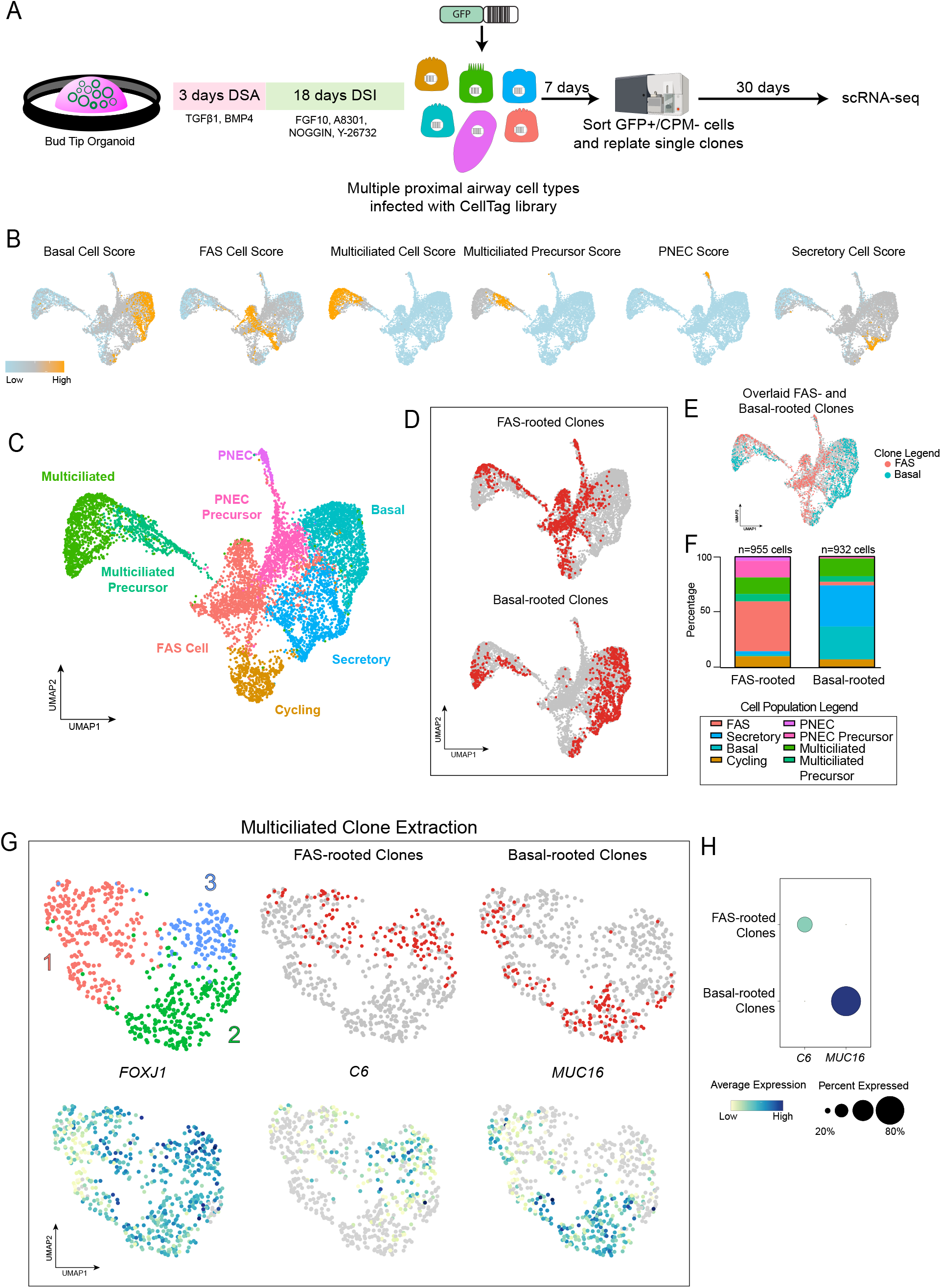
Barcoding-based lineage tracing identifies separate lineages from FAS cells and basal cells *in vitro*. (A) Schematic displaying experimental design for lineage-tracing BTOs using CellTagging. BTOs were differentiated using the 21-day Airway Differentiation Paradigm (Fig S2A). At 21 days, Airway Organoids were transduced with the CellTagging lentiviral-based library of barcoded GFP plasmids. After 7 days, cells were sorted for GFP^+^ and remaining BTPs were removed using CPM. Single clones were replated and grown for 30 days, then submitted for scRNA-sequencing. (B) UMAP feature plots of cell-type scoring annotation for predominant cell types on CellTagged-Airway Organoids. Cells most aligning to the scoring set are in orange, with little to no scoring represented in blue. (C) UMAP cluster plot of sequenced 58-day CellTagged-Airway Organoids. Each dot represents a single cell and cells were computational clustered based on transcriptional similarities. Clusters were colored and labeled by cell type, which were determined based on expression of ‘Cell Type Score’ (Fig. 2B, S2H, Table S2) derived from Fetal Epithelial dataset. (D) UMAP plots of CellTagged-Airway Organoids overlaid with cells labeled from FAS-Rooted or basal-rooted clones. (E) UMAP plot of FAS-rooted and basal-rooted clones overlaid. (F) Quantification of cell types within FAS- or basal-rooted clones. Number of cells per cell type were divided by total number of cells of -rooted clone sample to normalize and report as percentage. (G) UMAP cluster plot of multiciliated clone extraction from 58-day CellTagged-Airway Organoids. CellTag-expressing cells within the multiciliated cluster in Fig. 2C were computationally extracted and sub-clustered resulting in 3 subclusters. Cells from FAS-rooted or basal-rooted clones are shown as red dots in top middle and right panels. Bottom panels are UMAP feature plots of multiciliated marker *FOXJ1*, and enriched genes *C6* and *MUC16*. The color of each dot in the feature plot indicates log-normalized expression level of the genes in the represented cell. (H) Dot plot for expression of *C6* and *MUC16* between FAS-rooted and basal-rooted clones within multiciliated clone extraction (Fig. 2G). The dot size represents the percentage of cells expressing the gene in the corresponding cluster, and the dot color indicates log-normalized expression level of the gene.

Next, we investigated clone distributions to determine if there were stark trends or patterns associated among the clones. Clones that possessed between 2-20 unique tags were included in the analysis; using a correlation coefficient of 0.7 (2), resulting in 279 total unique clones identified within the dataset. We further filtered data to include clones containing ≥10 cells, resulting in 43 clones used in subsequent analysis. By interrogating individual clones, we began to observe a general trend that clones contained either tagged FAS cells or tagged basal cells but did not possess robust tagged populations of both cell types simultaneously (Figure 2D). Clones were then evaluated by their cellular composition and were specifically interrogated for the total number of FAS and basal cells they contained. Clones that contained ≥80% FAS cells were classified as “FAS-rooted”, and clones containing ≥80% basal cells were classified as “basal-rooted” (Figure 2D). Of the 43 clones analyzed, there were 24 FAS-rooted clones (n = 955 cells) and 13 basal-rooted clones (n = 932 cells). Among the FAS-rooted and basal-rooted clones, cells occupy distinct regions of the UMAP (Figure 2E) suggesting they represent distinct cell lineages. We quantified the distribution of cells attributed to FAS-rooted and basal-rooted clones (Figure 2F) and observed that even with the 80% threshold, a very small portion of FAS cells are present in basal-rooted clones and vice versa (all FAS-rooted clones are composed of 1.6% basal cells, while all basal-rooted clones contain 3.3% FAS cells). Striking trends are found among the remaining cell types: PNECs and the *ASCL1*^+^ PNEC precursors predominantly share CellTags with FAS-rooted clones (19% of cells in FAS-rooted clones are PNECs or PNEC precursors, compared to only 2% of cells in basal-rooted clones); secretory cells make up only 3% of cells in FAS-rooted clones compared to 37% of cells in basal-rooted clones. Finally, both populations contained a similar percentage of multiciliated and multiciliated precursor cells (21.8% of cells in FAS-rooted, 21% of cells in basal-rooted). Collectively, this data suggests lineages in the human airway bifurcate downstream of bud tip progenitors, with FAS cells sharing a cellular origin with the PNEC/PNEC precursor and multiciliated/multiciliated precursors, but not with basal or secretory cells, while basal cells share a cellular origin with secretory and multiciliated/multiciliated precursors but not with FAS cells or PNECs.

### FAS cells and basal cells contribute to multiciliated cell heterogeneity

While FAS-rooted clones and basal-rooted clones preferentially share CellTags with some lineages over others, the common cell population shared by both were multiciliated cells and multiciliated precursors. However, we noted that FAS and basal-rooted clones occupied distinct spatial domains within the multiciliated cell cluster, indicating they are transcriptionally distinct (Figure 2E). To explore transcriptional heterogeneity amongst multiciliated cells during *in vitro* airway differentiation, we computationally extracted and sub-clustered the multiciliated and multiciliated precursor clones from the organoid data (Figure 2G). Sub-clustering (45) identified 3 sub-clusters within the extracted multiciliated cells. Among these cells, the FAS-rooted and basal-rooted clones occupy distinct sub-clusters: FAS-rooted clones predominate cluster 3 (blue) and top subset of cluster 1 (pink); basal-rooted clones occupy cluster 2 (green) and bottom subset of cluster 1 (pink) (Figure 2G). Differential gene expression across clusters (Table S3) allowed us to identify two genes that distinguish FAS- or basal-rooted multiciliated cells. The complement component *C6* is enriched within the FAS-rooted subset while *MUC16*, which was recently identified as a multiciliated marker in the adult trachea (6), is specific to the basal-rooted clusters (Figure 2G-H).

As a result of the suggested lineage relationships from the CellTagging experiment, we proposed a revised model of airway differentiation within the developing human lung, with FAS and basal cells as separate progenitor cell populations differentiating from the common bud tip progenitor, giving rise to distinct differentiated cell types including transcriptionally distinct multiciliated cells (Figure 3A). To begin to test this model, and to validate the multiciliated cell heterogeneity observed *in vitro*, we investigated the multiciliated cell population in the developing human lung epithelium (Figure 3B). *C6* was indeed enriched in the multiciliated cluster while *MUC16* was seen in the multiciliated, multiciliated precursor, and club-like secretory clusters at lower expression levels (Figure 3B). Similar to the organoid data (Figure 2), within the multiciliated clusters, these genes quantitatively appeared to occupy separate regions. To interrogate this heterogeneity further, we computationally extracted the multiciliated cluster from the human epithelial data and sub-clustering predicted 5 sub-clusters (Figure 3C). All 5 clusters expressed canonical multiciliated marker *FOXJ1* at varying levels; other secretory markers were evaluated to define the subclusters given the expected secretory-to-multiciliated transition in both lineages (Figure 3C). Cluster 3 expressed the highest levels of *FOXJ1* but also had negligible levels of *C6* or *MUC16*, and expressed very low *TP63*, as well as secretory markers *SCGB1A1* and *SCGB3A2*, suggesting these are transitioning secretory cells. Clusters 1 and 2 on the other hand, were similar to multiciliated cells observed *in vitro*.

**Figure 3:**
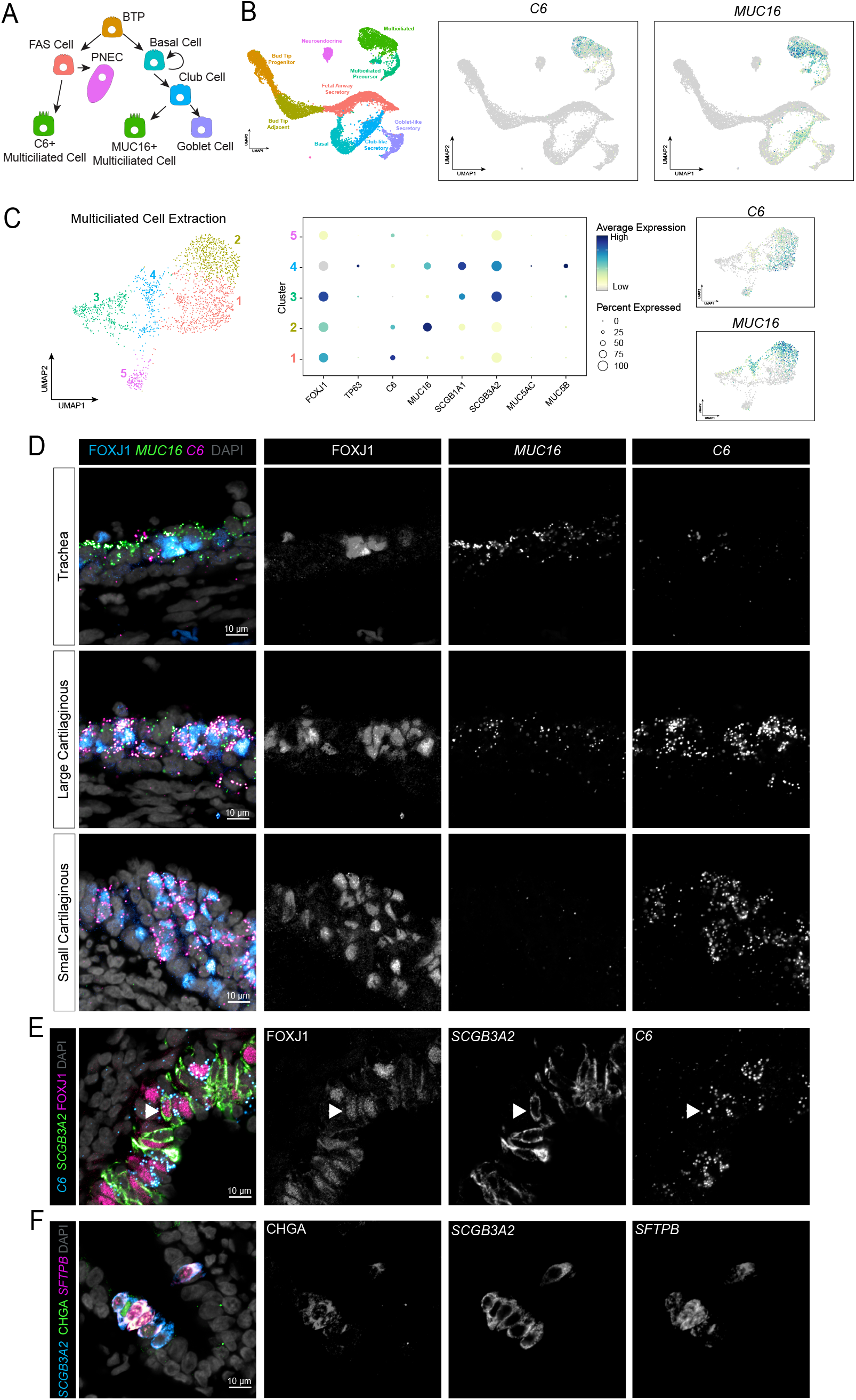
*C6* and *MUC16* distinguish subsets of multiciliated cells within developing human lung. (A) Hypothesized model of airway differentiation cellular hierarchy from CellTagging predictions. (B) UMAP feature plots of *C6* and *MUC16* on fetal lung epithelium UMAP from Fig 1A, reintroduced here. The color of each dot in the feature plot indicates log-normalized expression level of the genes in the represented cell. (C) UMAP cluster plot of multiciliated cell extraction from fetal epithelium data. The multiciliated cell cluster was computationally extracted and re-clustered resulting in 5 subclusters. The plot is colored and numbered by cluster. Middle panel is a dot plot for expression of various airway epithelial cell markers within multiciliated cell extraction. The dot size represents the percentage of cells expressing the gene in the corresponding cluster and the color indicates log-normalized expression level of the gene. Feature plots for *C6* and *MUC16* on multiciliated cell extraction plot. The color of each dot in the feature plot indicates log-normalized expression level of the genes in the represented cell. (D) FISH with co-IF staining on paraffin sections of 11.5-PCW fetal lung representative for multiciliated cell markers *C6, MUC16*, and FOXJ1, across different lung regions ranging from trachea, large cartilaginous, and small cartilaginous airways. (E) FISH with co-IF staining on paraffin sections of 12-PCW fetal lung representative for FAS-rooted multiciliated marker *C6*, alongside FAS marker *SCGB3A2*, and canonical multiciliated cell marker FOXJ1. Arrowheads highlight triple-expressing cell. (F) FISH with co-IF staining on paraffin sections of 12-PCW fetal lung representative for FAS markers *SCGB3A2* and *SFTPB*, with neuroendocrine cell marker CHGA showcasing triple-expressing cells.

Both clusters were *FOXJ1*^+^ with cluster 1 being *C6*^*+*^*/MUC16*^*-*^ and cluster 4 being *C6*^*LOW*^*/MUC16*^+^ (Figure 3C). We performed FISH for *C6* and *MUC16* with co-IF for FOXJ1 on 11-12 PCW human lung samples across the proximal-distal axis in regions containing multiciliated cells at this age (trachea, large cartilaginous, small cartilaginous airway) (Figure 3D). The regions examined also corresponded to areas of low and high abundance of basal and FAS cells, respectively, as demonstrated in Figure 1.

Interestingly, *MUC16* and *C6* expression was inversely correlated across the proximal-distal axis of the airway. That is, we observed that *MUC16*^*+*^ cells are localized predominantly in the trachea, while *C6* expression was very low in the trachea. As we interrogated large cartilaginous and small cartilaginous airways, *C6*^+^ cells become increasingly abundant while *MUC16* expression was reduced (Figure 3C).

Given the clonal relationship between FAS cells, multiciliated/multiciliated precursors and PNEC/PNEC precursors in the CellTagging data (Figure 2), we also examined primary lung tissue sections to determine if we could identify FAS cells in the process of differentiation toward either of these cell types (Figure 3E). We carried out FISH with co-IF for *C6/SCGB3A2*/FOXJ1 to identify cells with both FAS and multiciliated cell markers (Figure 3E) and *SCGB3A2/SFTPB/*CHGA to identify cells with both FAS and PNEC markers (Figure 3F). In both cases, cells co-expressing FAS cell markers with multiciliated cell (FOXJ1^+^) or PNEC (CHGA^+^) markers, respectively, were readily observed. Collectively, this data lends further support to our findings from the *in vitro* airway organoid differentiation by showing distinct subsets of multiciliated cells are anatomically correlated in abundance with the proposed cell of origin *in vivo* and that C6^+^ multiciliated cells and PNECs arise directly from FAS cells.

### Functional assessment of basal and FAS cell lineages *in vitro*

To further test the proposed lineage hierarchy (Figure 3A) we used FACS to isolate basal and FAS cells from DSA/DSI differentiated BTOs, grow as separate cultures, and then assess what cell types these organoids gave rise to. We predicted the FAS cultures would primarily give rise to *C6*^+^ multiciliated cells and PNECs, while basal cell cultures would give rise to *MUC16*^+^ multiciliated cells and secretory cells. To enrich FAS cells and basal cells, we used well-established surface markers for lung epithelial cell types and applied both a positive and negative selection sorting strategy (Figure 4A). BTPs were isolated using the surface antigen CPM (40, 41) and removed from further analysis; basal cells were sorted using a previously described strategy gating on F3 +/-EGFR (1), and the remaining CPM^-^/F3^-/^EGFR^-^ fraction was considered ‘FAS-enriched’ (Figure 4A, S3A). Basal and FAS-enriched cells were collected after sorting and Cytospin analysis was used on a fraction of collected cells to validate enrichment of cell types between the populations. TP63 immunofluorescence was used to identify basal cells, while SFTPB was used to identify FAS cells in Cytospins (Figure S3B). TP63 antibody expression was significantly increased in the basal sorted culture compared to unsorted or FAS-enriched cultures, while SFTPB was significantly lower in the basal sorted cells compared to unsorted or FAS-enriched (Figure S3C).

**Figure 4:**
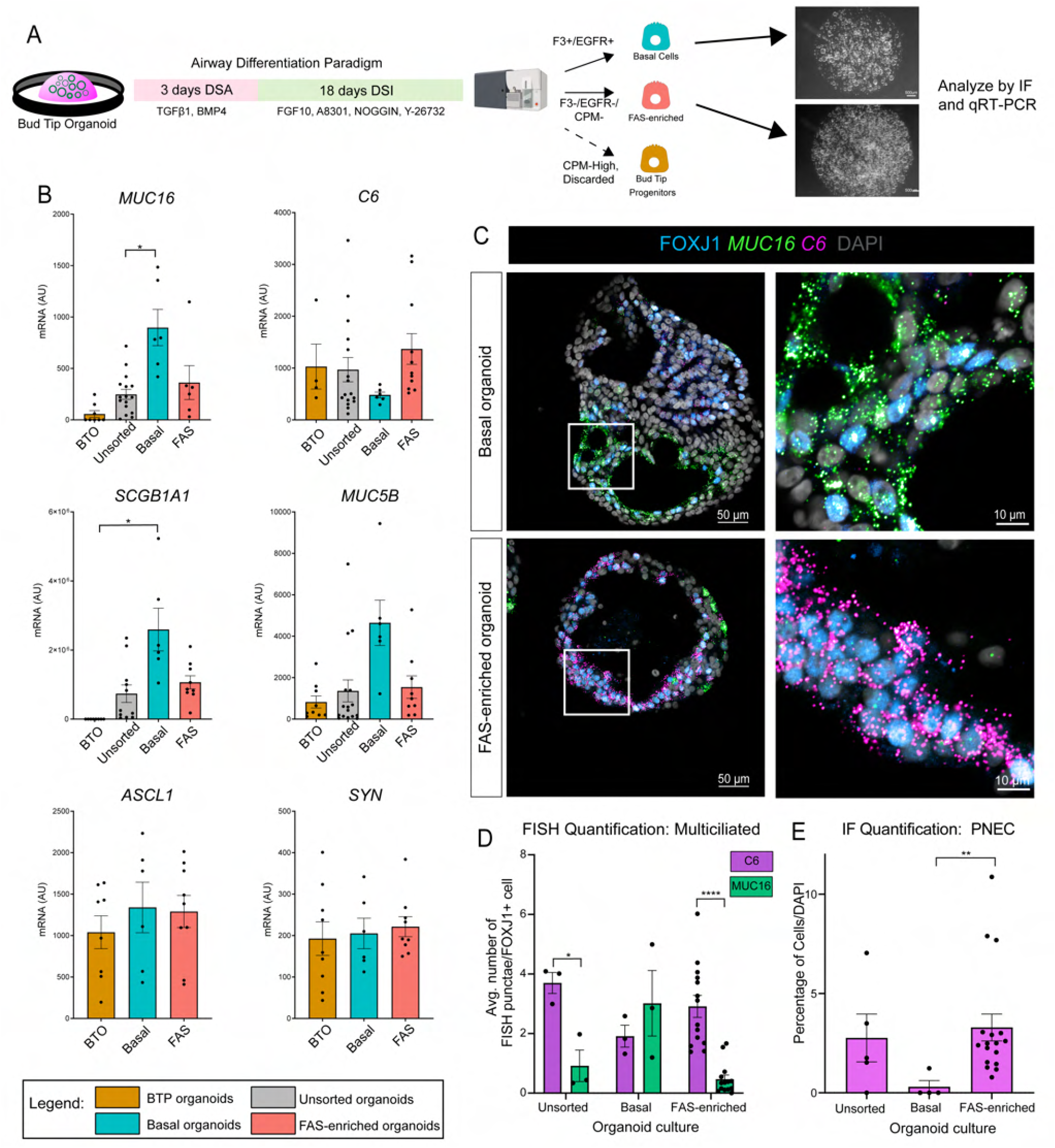
FAS-enriched cultures give rise to *C6*-multiciliated cells and PNECs. (A) Schematic of experimental design for separating basal and FAS cells *in vitro*. 21-day Airway Organoids are sorted using fluorescence activated cell sorting (FACS) by F3/EGFR for basal cells, CPM-High for BTPs (removed from culture) and triple-negative culture collected as ‘FAS-enriched’. Basal cell culture and FAS-enriched were replated to form organoids and analyzed after additional 2.5 weeks. (B) RT-qPCR data comparing expression of multiciliated (*MUC16, C6)*, secretory (*SCGB1A1, MUC5B)*, and PNEC (*ASCL1, SYN*) cells between bud tip progenitor organoids (BTO, orange), unsorted airway organoids (grey), basal organoids (teal) and FAS-enriched organoids (pink) 2.5 weeks post-sort. This quantification was performed on two to three unique biological replicates with at least three technical replicates. Error bars represent standard error of the mean. Statistical tests were performed by one-way ANOVA with Welch’s correction; p-values are (*) <0.05. (C) FISH and IF stains on paraffin sections of basal and FAS-enriched organoids collected 2.5 weeks post-sort for multiciliated cell markers (*C6, MUC16*, FOXJ1). (D) Quantification of multiciliated signal from FISH images for unsorted control, basal and FAS-enriched organoids. Total number of FISH punctae for *C6* or *MUC16* were averaged across FOXJ1^+^ cells in images. This quantification was performed in one to three unique biological replicates with at least three technical replicates (organoids per specimen). Error bars represent standard error of the mean. Statistical tests were performed by Welch’s t-test (unpaired, two-tailed) comparing C6 to MUC16 for each condition; p-values are (*) <0.05, (****) <0.0001, and 0.4252 (Basal). For comparison of each marker across conditions, one-way ANOVA with Welch’s correction was used; p-values >0.05 for all. (E) Quantification of neuroendocrine cell marker expression from IF images for unsorted control, basal and FAS-enriched organoids. Total number of ASCL1^+^ and/or CHGA^+^ cells were quantified and calculated as percentage of DAPI^+^ cells in image. This quantification was performed in one to three unique biological replicates with at least three technical replicates (organoids per specimen). Error bars represent standard error of the mean. Statistical tests were performed by one-way ANOVA with Welch’s correction; p-values are (**) <0.005, 0.2518 (Unsorted-basal), and 0.9716 (Unsorted-FAS).

Cells not used for Cytospin analysis were replated in 3D matrix, where cultures reformed cysts within 24 hours of sorting and were grown for an additional 2.5 weeks to allow for differentiation before collecting for qRT-PCR and IF (Figure 4A). qRT-PCR was performed to identify markers of expected cell lineages between basal and FAS-enriched organoids and compared to both BTOs and unsorted airway organoids (Figure 4B). In the basal-rooted organoids, *MUC16*, and secretory markers *SCGB1A1* and *MUC5B* were all increased compared to the other 3 organoid populations, while in the FAS-enriched organoids, *C6* was increased compared to basal organoids (Figures 4B). PNEC markers (*ASCL1, SYN*) were consistent across BTOs, basal and FAS-enriched organoids (Figure 4B), and significantly lower than the unsorted organoids (Figure S3D). Other epithelial markers were unchanged between the cultures (*SOX2, SOX9*) (Figure S3D). Organoid sections were also stained for multiciliated markers (*C6/MUC16/*FOXJ1), PNECs (ASCL1/CHGA), FAS markers (*SCGB3A2/SFTPB/CFTR*), bud tips (SOX9) and basal cells (TP63) (Figures 4C, S3E). Images were quantified for cell types present, which show similar trends as the qRT-PCR data (Figures 4D-E, S3F-G). FAS-enriched organoids contain significantly more *C6*^+^/FOXJ1^+^ cells while basal organoids contain more *MUC16*^+^/FOXJ1^+^ cells (Figure 4C-D). Despite the low level of PNEC marker expression by bulk qRT-PCR, IF of organoid cultures revealed a small population of PNECs (ASCL1^+^ or CHGA^+^) that was significantly higher in FAS-enriched organoids compared to basal organoids (Figure 4E, S3E arrowheads). Additionally, FAS-enriched organoids contain higher numbers of FAS cells compared to basal organoids (Figure S3F), while the basal organoids have more TP63^+^ cells (Figure S3G). Neither basal nor FAS-enriched organoids contain many SOX9^+^ cells. suggesting bonafide bud tips are not present in culture at this timepoint (Figure S3E, S3G).

As an alternative and independent method to interrogate the model proposed in Figure 3A, fetal tracheal epithelium was isolated directly from primary tissue and basal cells were expanded using a conditional reprogramming culture method (see ‘Methods’). Following expansion, basal cells were seeded into 3D organoid culture or onto 2D air-liquid interface culture, differentiated *in vitro* in DSI media, and both conditions were subsequently analyzed by scRNA-seq (see ‘Methods’; Figure S4A-B). We observed that *FOXJ1*^+^ multiciliated cells were present within the cultures and expressed robust *MUC16* (Figure S4B-C), but low levels of *C6*, similar to observations made in multiciliated cells from primary lung epithelium (Figure 3B-C). Notably, no PNEC markers were detected in the dataset and there were no cells expressing the FAS gene signature *SCGB3A2*^*+*^*/SFTPB*^*+*^*/CFTR*^*+*^*/SCGB1A1*^*-*^ (Figure S4B). FISH and IF carried out on these organoids supported the trends shown by scRNA-seq, with a majority of FOXJ1^+^ cells expressing *MUC16* but very little *C6*; rare ASCL1^+^ or CHGA^+^ cells were detected by IF and no FAS-like cells were detected (Figure S4D). These data suggest that primary tracheal organoids derived from tracheal basal cells give rise to MUC16^HI^/C6^LO^ multiciliated cells, but rarely give rise to PNECs and do not give rise to FAS cells. Taken together, this analysis demonstrates that enrichment of organoid cultures for either FAS or basal cells also enriches for cell types predicted as part of the FAS or basal lineage CellTagging. More specifically, these findings support our proposed lineage hierarchy, where FAS cells and basal cells bifurcate downstream of BTPs to give rise to specific cell types as well as transcriptionally distinct multiciliated cell subtypes during human lung development.

## DISCUSSION

With widespread use of single cell genomic technologies, the field has been able to study cellular diversity of the human lung at an unprecedented level, leading to the identification of new cell types and cell states (1, 3, 6, 8, 10, 13–18, 20, 21, 23–27). Functional assessment of these new cell types and states is essential to understand how these cells contribute to the unique physiology of the lung. Given the uniquely human biology being uncovered, animal models are not always appropriate for functional follow-up studies, increasing our reliance on *in vitro* human models. Here, we characterize a previously unreported secretory cell (FAS cells) within the developing human lung defined by the co-expression of *SCGB3A2/SFTPB/CFTR*. Using *in situ* analysis on tissue sections and an *in vitro* organoid model system coupled with single cell-barcoding lineage tracing techniques, we demonstrate that FAS cells are a distinct progenitor cell in the airway that give rise to pulmonary neuroendocrine cells and a subset of multiciliated cells within the lung marked by the expression of complement component *C6*, while the basal cell lineage gives rise to multiciliated cells defined by *MUC16* as well as secretory cells.

The heterogeneity of multiciliated cells has recently been explored in adult lungs, which also defines a *MUC16*^+^ subpopulation in the trachea (6). Our work demonstrates that this heterogeneity within cell types do not necessarily represent different states of the same cell, but similar cells of different origin. Additionally, an interesting finding from the current work is that basal cells are enriched in the trachea and cartilaginous airways while FAS cells are enriched in the small cartilaginous and non-cartilaginous airways. Moreover, it is well established that cells in the trachea are specified early during development while the lower regions of the airway are formed later during branching morphogenesis (46). It is interesting to speculate that in the human lung, basal cells and FAS cells are specified at different developmental times and are responsible for tuning the distinct cell type composition of airways along the proximal-distal axis. Consistent with this idea, large animal studies as well as studies in humans have defined functionally distinct subsets of multiciliated cells in different anatomical regions of the lung that have different ciliary beat frequencies (47–49). It is also possible that the changing cellular landscape of the airway epithelium along the proximal to distal axis provides an evolving defense mechanism, with a strong immune response (notably *C6*^+^ cells) becoming enriched before reaching the alveoli. Additional functional assessments of basal and FAS-derived multiciliated cell subtypes in the future will help to understand the importance of this heterogeneity.

The cell of origin for pulmonary neuroendocrine cells (PNECs) also remains unclear, with evidence supporting both bud tip progenitors and basal cells giving rise to PNECs (1, 42, 44, 50–54). It is possible that PNECs are derived from both cell lineages but in different contexts (development versus injury repair). Furthermore, recent data suggests there may be additional PNEC heterogeneity that has yet to be functionally resolved (14). PNECs are believed to be the earliest specified cells in airway development (55), and it is also possible that our findings are capturing this earliest stage of PNEC differentiation during development, while basal cell derived PNECs are more representative of a homeostatic adult lung. Given the importance of NOTCH signaling in both multiciliated and PNEC differentiation in the mouse lung (56–61), it will be interesting to carry out follow-up studies to understand how FAS-derived PNEC cells are regulated. Further understanding of PNECs within the human lung shed light on this unique cell type and its origin.

While the existence of a FAS cell has not been demonstrated in mice, it is likely that there are different airway progenitors that populate the lungs during development. It is well accepted that basal cells act as a stem/progenitor cell aiding in homeostasis and injury repair in mice; however, there is also evidence that basal cells are not required for airway development. For example, in the lungs of *P63*-KO mice, which lack basal cells and die shortly after birth, both secretory and multiciliated cells are still present during development, supporting the hypothesis that non-basal cells can populate the airway (52, 62, 63). The presence of cells expressing *Scgb3a2*+ have also been identified during murine lung development (30, 64). It is not clear whether these cells are analogous to the FAS cells, and future work comparing mouse and human lung development will help answer this question; however, existing evidence suggests functional differences. For example, *Scgb3a2+* cells in mice predominantly give rise to club and multiciliated cells, but not the PNEC lineage (64), while our results suggest that FAS cells give rise to the PNEC lineage, but not the club lineage.

The developmental potential of the FAS cell also warrants further investigation. The presence of *SCGB3A2*^+^*/SFTPB*^+^*/SCGB1A1*^-^ cells in the adult terminal respiratory bronchiole has recently been characterized, referred to as Terminal Respiratory Bronchiolar Stem Cells (TRB-SC) or Respiratory Airway Secretory Cells (RASC) (10, 24). It is currently unclear if FAS cells are the fetal equivalent to this cell, however both TRB-SC and RASCs were shown to give rise to alveolar cell types, while FAS cells give rise to airway cell types, at least under the conditions tested here. The differences, similarity, and relationship between these populations across the lifespan remains unclear until more data can be acquired later in development and early in postnatal life.

Taken together, the current work identifies FAS cells as an important cell type during human lung development which has broadened our understanding of airway epithelium and challenged how we approach differentiation in this organ. As new techniques and technologies further our exploration of tissue heterogeneity, more cell types will emerge that warrant investigation and functional evaluation in all organs.

## MATERIALS AND METHODS

### Human Lung Tissue

Human lung tissue research was reviewed and approved by The University of Michigan Institutional Review Board (IRB). Normal, de-identified human lung tissue was obtained from the University of Michigan Laboratory of Developmental Biology. Tissue was shipped overnight in UW-Belzer’s solution (Thermo Fisher, NC0952695) on ice and was processed for experiments or fixation within 24h.

### Cell Lines & Culture Conditions

#### BTO Line Establishment and Culture

BTO lines were cultured as previously described (1, 29). Briefly, distal lung tissue was dissected into 1 cm^3^ chunks and washed with sterile PBS. Tissue was digested using 1 mL Dispase (Corning, Cat#354235) for 30 min on ice. Digestion was then quenched with 1 mL 100% FBS (Thermo Fisher, Cat#35-015-CV) for 15 min on ice following by washing in 1 mL media containing DMEM-F12, 1% Penicillin-streptomycin (Thermo Fisher, Cat#15140122) and 10% FBS for 10 min on ice. Tissue was then mechanically dissociated further using a scalpel, forceps, and vigorous pipetting with a P1000 to remove mesenchyme from bud tip epithelium. Dissociated tissue was transferred to a 15 mL conical tube, washed with DMEM-F12 media described above and spun at 300g for 3 min at 4°C. After each wash, supernatant and separated mesenchyme was removed and fresh DMEM-F12 was added. These steps were repeated 3-5 times until all visible mesenchyme is removed. On final wash, remaining epithelium was transferred to a 1.5 mL microcentrifuge tube and cells were resuspended in Matrigel (Corning, Cat#354234). Cells were plated into 15 µL droplets and allowed to solidify. BTOs were fed 3F media (29) every 3 days and were passaged using 27G needle every 10-14 days.

#### Airway Differentiation Paradigm

Airway differentiation was carried out as previously described (1). Briefly, BTOs were exposed to dual-SMAD activation (DSA) via 100ng/mL BMP4 (R&D Systems, Cat#314-BP-050) and 100ng/mL TGFβ1 (R&D Systems, Cat#240-B-002) in 3F media for 3 days. On the fourth day, BTOs were exposed to dual-SMAD inactivation (DSI) via 1µM A8301 (APExBIO, Cat#3133), 100ng/mL NOGGIN (R&D Systems, Cat#6057), 10µM Y-27632 (APExBIO, Cat#B1293) and 500ng/mL FGF10 (lab purified – see below) in serum-free basal media for 18 days (media changed every 3 – 4 days) with needle-passaging as needed.

#### Tracheal Basal Cell Airway Cultures Establishment, Culture, and Single-Cell Dissociation

Proximal tracheal tissue was dissected from tissue and epithelial cells were scraped from surface using a scalpel. Epithelial tissue was seeded onto 2D culture with irradiated 3T3-J2’s where basal cells expanded in DSI media. After 2D expansion, cultures were seeded in 3D and air-liquid interface (ALI) cultures. Briefly, cells were spun down at 300g for 3 min, supernatant media was removed, and cells were resuspended in Matrigel. Cells were replated in 20 µL droplets and allowed to solidify. Airway organoids (AOs) were fed 0.5 mL DSI media every 3-4 days. Cultures were grown for 40 days before collection for scRNA-seq. The Neural Tissue Dissociation Kit (P) (Miltenyi, Cat#130-092-628) was used for single cell dissociation of 3D cultures. For ALI cultures, cells were seeded onto Costar transwells (Corning, Cat#3470). ALI cultures were fed 0.5 mL DSI media every 3-4 days. 5 days after seeding, cultures were exposed to air, and then grown for an additional 14 days before collection for scRNAseq. Single cell dissociation of ALI cultures included a 2 min exposure to 0.25% Trypsin at 37°C (Gibco, Cat#25200-056), careful aspiration of Trypsin, followed by 5 min of Accutase (Sigma, Cat#A6964) incubation at room temperature.

#### Expression and Purification of Human Recombinant FGF10

The recombinant human FGF10 (rhFGF10) expression plasmid pET21d-FGF10 was a gift from James A. Bassuk (65)at the University of Washington School of Medicine. This plasmid was transformed to Novagen’s Rosetta™ 2(DE3)pLysS competent cells (Millipore Sigma, Cat#71403-3) for rhFGF10 expression. In brief, *E*.*coli* strain Rosetta™ 2(DE3)pLysS bearing pET21d-FGF10 was grown in 2x YT medium (BD Biosciences, Cat#244020) with Carbenicillin (50µg/ml) and Chloramphenicol (17µg/ml). rhFGF10 expression was induced by addition of isopropyl-1-thio-β-D-galactopyranoside (IPTG). rhFGF10 was purified by using a HiTrap-Heparin HP column (GE Healthcare, Cat#17040601) with step gradients of 0.2M to 0.92M NaCl. From a 200mL culture, 3 – 4mg of at least 98% pure rhFGF-10 (evaluated by SDS-PAGE stained with Coomassie Blue R-250) was purified. In-house purified rhFGF10 was confirmed by western blot analysis using anti-FGF10 antibody and compared to commercially purchased rhFGF10 (R&D Systems, Cat#345-FG) to test/validate activity based on the efficiency to phosphorylate ERK1/2 in an A549 alveolar epithelial cell line (ATCC, Cat#CCL-185) as assessed by western blot analysis

### Tissue processing, Staining, and Quantification

All sectioned fluorescent images were taken using a Nikon A1 confocal microscope, an Olympus IX83 inverted fluorescence microscope or an Olympus IX71 inverted fluorescence microscope. Acquisition parameters were kept consistent for images in the same experiment and all post-image processing was performed equally on all images in the same experiment. Images were assembled in Adobe Photoshop CC 2022.

#### Tissue Processing

Tissue was immediately fixed in 10% Neutral Buffered Formalin (NBF) for 24h at room temperature on a rocker. Tissue was then washed 3x in UltraPure DNase/RNase-Free Distilled Water (Thermo Fisher, Cat#10977015) for 15 min each and then dehydrated in an alcohol series of concentrations dehydrated in UltraPure DNase/RNase-Free Distilled Water for 1h per solution: 25% Methanol, 50% Methanol, 75% Methanol, 100% Methanol, 100% Ethanol, 70% Ethanol. Dehydrated tissue was then processed into paraffin blocks in an automated tissue processor (Leica ASP300) with 1 hr solution changes. 5 (FISH) or 7 (IF) µm-thick sections were cut from paraffin blocks onto charged glass slides. For FISH, microtome and slides were sprayed with RNase Away (Thermo Fisher, Cat#700511) prior to sectioning (within one week of performing FISH). Slides were baked for 1 hr in 60°C dry oven (within 24 hr of performing FISH). Slides were stored at room temperature in a slide box containing a silica desiccator packet and the seams sealed with paraffin.

#### Immunofluorescence (IF) Protein Staining on Paraffin Sections

Tissue slides were rehydrated in Histo-Clear II (National Diagnostics, Cat#HS-202) 2x for 5 min each, followed by serial rinses through the following solutions 2x for 3 min each: 100% EtOH, 95% EtOH, 70% EtOH, 30%EtOH, and finally in double-distilled water (ddH2O) 2x for 5 min each. Antigen retrieval was performed using 1X Sodium Citrate Buffer (100mM trisodium citrate (Sigma, Cat#S1804), 0.5% Tween 20 (Thermo Fisher, Cat#BP337), pH 6.0), steaming the slides for 20 min, followed by cooling and washing quickly 2x in ddH2O and 2x in 1X PBS. Slides were incubated in a humidified chamber at RT for 1 hr with blocking solution (5% normal donkey serum (Sigma, Cat#D9663) in PBS with 0.1% Tween 20). Slides were then incubated in primary antibody diluted in blocking solution at 4°C overnight in a humidified chamber. Next, slides were washed 3x in 1X PBS for 5 min each and incubated with secondary antibody with DAPI (1µg/mL) diluted in blocking solution for 1h at RT in a humidified chamber. Slides were then washed 3x in 1X PBS for 5 min each and mounted with ProLong Gold (Thermo Fisher, Cat#P36930) and imaged within 2 weeks. Stained slides were stored in the dark at 4°C. All primary antibody concentrations are listed in Supplementary Table 4. Secondary antibodies, raised in donkey, were purchased from Jackson Immuno and used at a dilution of 1:500.

#### Fluorescence in situ hybridization (FISH)

The FISH protocol was performed according to the manufacturer’s instructions (ACDbio, RNAscope multiplex fluorescent manual) with a 5-minute protease treatment and 15-minute antigen retrieval. For IF co-staining with antibodies, the last step of the FISH protocol was skipped and instead the slides were washed 1x in PBS followed by the IF protocol above. A list of probes and reagents can be found in Supplementary Table 4.

IF and FISH stains were repeated on at least 3 independent experiments and representative images are show.

### Organoid Tissue Prep for scRNA-seq

All tubes and pipette tips were pre-washed in 1X HBSS with 1% BSA to prevent cell adhesion to the plastic. Organoid cultures were removed from Matrigel using a P1000 pipette tip and vigorously pipetted in a 1.5mL microcentrifuge tube to remove as much Matrigel as possible. Tissue was centrifuged at 300g for 3 min at 4°C, then excess media and Matrigel was removed. Tissue was digested to single cells using 0.5mL TrypLE (Invitrogen, Cat#12605010) and incubated at 37°C for 30 min, adding mechanical digestion with pipette every 10 min. After 30 min, trypsinization was quenched with 1X HBSS. Cells were passed through a 40µm filter (Bel-Art Flowmi, Cat#136800040), and centrifuged at 300g for 3 min at 4°C. Cells were resuspended in 1mL 1X HBSS and counted using a hemocytometer, centrifuged at 300g for 3 min at 4°C and resuspended to a final concentration of 1,100 cells/µL. Approximately 100,000 cells were put on ice and single cell libraries were immediately prepared at the 10X Chromium at the University of Michigan Sequencing Core with a target of 10,000 cells per sample.

### RNA extraction, cDNA, qRT-PCR

Each analysis includes three technical replicates from three separate biologic tissue lines. mRNA was isolated using the MagMAX-96 Total RNA Isolation Kit (Thermo Fisher, Cat#AM1830). RNA quality and yield was measured on a Nanodrop 2000 spectrophotometer just prior to cDNA synthesis. cDNA synthesis was performed using 100ng RNA per sample with the SuperScript VILO cDNA Kit (Thermo Fisher, Cat#11754250). qRT-PCR was performed on a Step One Plus Real-Time PCR System (Thermo Fisher, Cat#42765592R) using QuantiTect SYBR Green PCR Kit (Qiagen, Cat#204145). Primer sequences can be found in Supplementary Table 4. Gene expression as a measure of arbitrary units was calculated relative to Housekeeping gene (GADPH, ECAD or FOXJ1 for multiciliated markers) using the following equation:

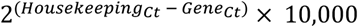

### Quantification, Statistical Analysis

Quantification of immunofluorescence images was performed using CellProfiler (66, 67) or by a blind count method. All images for FISH quantification were taken at 40X magnification, IF images were taken at 20X or 40X magnification. Briefly, CellProfiler pipelines were established to perform quantification of number of speckles for *C6* or *MUC16* FISH expression within a masked region for FOXJ1 for multiciliated counts. Nuclear stains were counted by CellProfiler to count nuclei, TP63^+^ cells for basal count, SOX9^+^ cells for BTP count. ASCL1^+^ cells were counted by hand for PNEC count in blinded fashion by independent researcher. FAS cells were counted by hand for triple-expressing cells in blinded fashion by independent researcher. For all quantification of images, at least 3 independent organoids were counted (technical replicates) from n=2-3 independent biological specimens (biological replicates) among at least n=3 experimental replicates. For figure 1H, 100 DAPI^+^ cells within the epithelium were counted for each airway region per sample and results were plotted as percentage of SCGB3A2^+^/SFTPB^+^ cells per 100 DAPI^+^ cells. For qRT-PCR analysis, n=2-3 biological replicates were used. For each biologic replicate, 3 wells of organoids containing 10-60 organoids (technical replicates) were collected.

All statistical analysis (graphs and statistical analysis for RT-qPCR and IF image quantification) were performed in GraphPad Prism Software. See Figure Legends for the number of replicates used, statistical test performed, and the p-values used to determine the significance for each separate analysis.

### Cytospin

25% of cells from each group for FACS (unsorted, basal culture, FAS-enriched culture) were reserved to evaluate the percentage of basal or FAS cells within each culture condition. 200 µL of cell suspension was isolated in a separate 1.5 mL microcentrifuge tube and diluted with FBS to final concentration of 5% vol/vol. Cell suspensions were placed in clean cytospin cones and spun at 600g for 5 min on an Epredia Cytospin 4 (Fisher Scientific Cat#A78300003) to charged glass slides. Slides air dried for 5 min before fixation with 100% ice cold methanol for 10 min. Slides were air dried for 10 min, followed by 2x washes in PBS for 5 min each at RT on a rocker. Immunofluorescence staining was performed as mentioned above.

### CellTagging Library Prep and Lentiviral Transduction into Organoids

#### CellTagging Library Preparation

The CellTag V1 plasmid library was purchased from Addgene (‘CellTag A’, Cat#124591). 25 ng of plasmid library was transformed into 100 µl of Stellar Competent *E. coli* (Takara Biosciences, Cat#636763) at an efficiency of 1.79×10^9^ cfus/µg. Plasmid library was isolated from 500 mL transformed *E. Coli* using the Plasmid Plus Mega Kit (Qiagen, Cat#12981). To assess library complexity, PCR product from the library was submitted for high throughput DNA sequencing (MiSeq, Illumina), resulting in 13963 unique tags in the 90^th^ percentile for frequency. Lentiviral packaging was performed by the University of Michigan Vector Core.

#### Lentiviral Transduction of Organoids

CellTag V1 lentivirus was thawed on ice and combined 1:1 with DSI media. 21-day differentiated BTOs (Airway Organoids) were collected for transduction by dislodging Matrigel droplet with P1000 and transferring organoids into a 1.5 mL microcentrifuge tube. Samples were needle passaged by 2 passes through a 27G needle and spun down at 300g for 3 min at 4°C. Cells were resuspended in 1 mL of DSI media + 6µg/mL of polybrene and incubated at 37°C for 15 min with agitation every 2 minutes. Cells were again spun down at 300g for 3 min at 4°C and resuspended in 300 µL virus + DSI. Cells were incubated at 37C for 6 hr with agitation every hour. Following transduction, cells were spun down again at 300g for 3 min, resuspended in Matrigel at 1:4 dilution, and fed with 0.5 mL DSI media.

#### Limited Dilution Plating following FACS

Probability of cultures seeded after FACS to contain cells from the same clone were estimated to be 10% by the probability of coincidences function in R and based an approximation of the numbers of clones generated during viral transduction as well as the number of cells seeded in the sequenced culture.

### Fluorescent Activated Cell Sorting (FACS) and Flow Cytometry

Organoid cultures were removed from Matrigel using a P1000 pipette tip and vigorously pipetted in a 15mL conical tube to remove as much Matrigel as possible. Tissue was centrifuged at 300g for 3 min at 4°C, then excess media and Matrigel was removed. Tissue was digested to single cells using 2 – 4mL TrypLE (Invitrogen, Cat#12605010), depending on pellet size, and incubated at 37°C for 30 min, adding mechanical digestion with a pipette every 10 min. After 30 min, trypsinization was quenched with DMEM/F-12 (Corning, Cat#10-092-CV) + 10µM Y-27632 (APExBIO, Cat#B1293). Cells were passed through a 70µm cell strainer, pre-coated with DMEM/F-12 +10µM Y-27632 and centrifuged at 500g for 5 min at 4°C. Cells were resuspended in 4mL FACS Buffer (2% BSA, 10µM Y-27632, 100U/mL penicillin-streptomycin) and transferred into 5 mL FACS tubes (Corning, Cat#352063). Cells were centrifuged again at 300g for 3 min at 4°C, then resuspended in 1mL FACS buffer and counted. 10^5^ cells were placed into new FACS tubes for all controls (no antibody, DAPI only, individual antibodies/fluorophores) and all remaining cells were centrifuged and resuspended in FACS buffer for a concentration of 10^6^ cells/100µL. Primary antibodies were incubated for 30 min on ice. 3mL FACS buffer was added to each tube after 30 min and tubes were centrifuged at 300g for 3 min at 4°C. Cells were washed again with 3mL FACS buffer and centrifuged at 300g for 3 min at 4°C. Secondary antibodies were incubated for 30 min on ice.

Conjugated antibodies were incubated for 10 min on ice. 3mL FACS buffer was added to each tube after 30 min and tubes were centrifuged at 300g for 3 min at 4°C. Cells were washed again with 3mL FACS buffer and centrifuged at 300g for 3 min at 4°C. Cells were resuspended in FACS buffer and 0.2µg/mL DAPI was added to respective tubes. FACS was performed using a Sony MA900 cell sorter and accompanying software. Cells were collected in 1mL 3F media +10µM Y-27632. All primary and conjugated antibody concentrations are listed in Supplementary Table 4. Secondary antibodies were purchased from Jackson Immuno and used at a dilution of 1:500.

### Bioinformatics/scRNAseq

#### Quality Control

To ensure quality of the data, all samples were filtered to remove cells expressing too few or too many genes (Fig. 1A-F/Fig. 3B-C/Fig. S1A-E/Fig. S2H– <500, >12000; Fig. 2B-H/Fig. S2G-I – <500, >10000; Fig. S4D-F – <200, >12000,), with too low or too high UMI counts (Fig. S4A-C– <200, >120000), or a fraction of mitochondrial genes greater than 0.1. Following the above steps, a total of (Fig. 1A-B/Fig. 3B/Fig. S1A-D – 10614 cells, 24390 genes; Fig. 1C-F/ Fig. S1C,E – 721 cells, 24390 genes; Fig. 2B-F/Fig. S2G-I – 8256 cells, 24402 genes; Fig. 2G-H – 642 cells, 24402 genes; Fig. 3C – 1955 cells, 24390 genes; Fig. S4A-B – 5832 cells, 22500 genes; Fig. S4C – 292 cells, 22500 genes) were kept for downstream analysis and visualization.

#### Pre-Processing and Integration

After quality control, the standard pre-processing workflow includes normalization, scaling, and selection of highly variable features. On the other hand, Seurat’s SCTransform method also allows efficient pre-processing, normalization, and variance stabilization of molecular count data from scRNA-seq samples. Running this algorithm will reveal a model of technical noise in the scRNA-seq data through “regularized negative binomial regression”, whose residuals are returned as the SCTransform-normalized values that can be used for further downstream analysis such as dimension reduction. During the SCTransform process, we also chose to regress out a confounding source of variation – mitochondrial mapping percentage. When dealing with one sample or when the batch effect is not prominent, we utilized either standard pre-processing (Fig. 2B-F/ Fig. S2G-I) or SCTransform (Fig. 2G-H/ Fig. S4C) based on their individual performance. If evidence of batch effect was present, we chose to follow Seurat’s integration workflow due to its optimal efficiency in harmonizing large datasets. The three integration methods used are integration on LogNormalized datasets using reciprocal PCA (Fig. 1A-B/ Fig. 3B/Fig. S1A-B,D/Fig. S2G-I), integration on SCTransform-normalized datasets (Fig. S4A-B), and fastMNN (Fig. 1C-F/ Fig. 3C/ Fig. S1C,E). After completion of such batch correction, the influence of batch specific technical artifacts on clustering is reduced.

#### Dimension Reduction and Clustering

Principal component analysis (PCA) was conducted on the corrected expression matrix followed. Using the top principal components (Fig. 1A-B/Fig. 3B/Fig. S1A-B,D-E/S2H –12 principle components, Fig. 1C-F/ Fig. 2G-H/ Fig. 3C/ Fig. S1C,E/ Fig. S4A-B – 30 principal components, Fig. 2B-F/Fig. S2G,I – 10 principle components), a neighborhood graph was calculated for the 30 nearest neighbors. The UMAP algorithm was then applied for visualization on 2 dimensions. Using the Leiden algorithm, clusters were identified with a resolution of (Fig. 1C-F/ Fig. 2G-H/ Fig. 3C/ Fig. S1C,E/ Fig. S4A-C – 0.3). Using the Louvain algorithm, clusters were identified with a resolution of (Fig. 1A-B/Fig. 3B/Fig. S1A-B,D-E – 0.2; Fig. 2B-F/Fig. S2G,I – 0.3; Fig. 2G-H/Fig. S2C-E/Fig. S3A – 0.08).

#### Cluster Annotation and Cell Scoring

Using canonically expressed gene markers, each cluster’s general cell identity was annotated. Gene lists for cell scoring (Fig. 2B) are found in Supplemental Table 2 and application of cell scoring strategy is as previously described (61, 62). Briefly, cells were scored based on expression of a set of 50 marker genes per cell type. Gene lists were compiled by analyzing previously published data from human fetal lung epithelium (62, 63). The top 50 genes from each cell type were merged to create the gene sets for cell scoring. After obtaining the scaled expression values for the data set, scores for each cell were calculated with the AddModuleScore function of Seurat. Cell scores were visualized by feature plots.

#### Normalization for Visualization and Differential Gene Expression

As recommended by Seurat developers, we employed the method of log normalization on the standard RNA assay for graphing dot plots, feature plots, and conducting DGEs. Expression matrix read counts per cell were normalized by the total expression, multiplied by a scale factor of 10000, and finally log-transformed. For the differential gene expression testing, we only tested features that are first, detected in a minimum fraction of 0.25 in either of the two cell populations, and second, show at least 0.25-fold difference in log-scale between the two cell populations on average.

#### Analysis of CellTagged Airway Organoids

Using the CellTagR package (https://github.com/morris-lab/CellTagR) CellTags were extracted from processed single cell RNA-sequencing BAM files to generate a matrix of cell barcodes, unique molecular identifiers and CellTags. This matrix was then filtered for cell barcodes corresponding to cells as determined by the CellRanger pipeline and then subjected to CellTag sequencing error correction using Starcode (https://github.com/gui11aume/starcode). Whitelisting was performed to remove tags not detected during assessment of CellTag library complexity (see CellTag Library Preparation). For clone calling, cells expressing less than 2 unique or more than 20 unique CellTags were removed and Jaccard analysis (Jaccard, 1912) was performed to calculate pairwise similarity coefficients between combinations of cell tags. Cells with CellTag combinations having similarity scores better than 0.7 were called as clones. For scRNA-seq analysis and visualization the standard Seurat workflow was performed including cell filtering by number of features (<500, >10000 removed) and percentage of mitochondrial reads (>10% removed), normalization, variable feature selection (n = 500), dimensional reduction (10 principle components) and Louvain clustering (resolution = 0.3). Clonal identities were appended as metadata in a Seurat object by cell barcode using Seurat’s AddMetaData function. To ensure clones were sufficiently sampled we limited our lineage analysis to clones containing ≥10 cells.

## ACKNOWLEDGEMENTS & FUNDING

We would like to thank Judy Opp and the University of Michigan Advanced Genomics core for the operation of the 10X Chromium single cell capture platform, the Microscopy core for providing access to confocal microscopes, the Viral Vector core for CellTagging lentivirus library prep, and the Flow Cytometry core for providing access to flow cytometers. We would also like to thank the University of Washington Laboratory of Developmental Biology and Beth Moore’s lab for use of their Cytospin.

This work was supported in part by grant CZF2019-002440 from the Chan Zuckerberg Initiative DAF, an advised fund of the Silicon Valley Community Foundation, and by the National Heart, Lung, and Blood Institute (NHLBI; R01HL119215) to JRS. ASC is supported by the T32 Michigan Medical Scientist Training Program (5T32GM007863-40) and by a Ruth L. Kirschstein Predoctoral Individual National Research Service Award (NIH-NHLBI F30HL156474). TF is supported by a NIH Tissue Engineering and Regenerative Medicine Training Grant (NIH-NIDCR T32DE007057). PPH is supported by the Rogel Cancer Center Fellowship, Judith Tam ALK NSCLC Research Initiative. A.J.M. was supported by a Ruth L. Kirschstein Predoctoral Individual National Research Service Award (NIH-NHLBI F31HL142197). RFCH is supported by a NIH Tissue Engineering and Regenerative Medicine Training Grant (NIH-NIDCR T32DE007057) and by a Ruth L. Kirschstein Predoctoral Individual National Research Service Award (NIH-NHLBI F31HL152531). EMH was supported by a Ruth L. Kirschstein Predoctoral Individual National Research Service Award (NIH-NHBLI F31HL146162). I.G. and the University of Washington Laboratory of Developmental Biology was supported by the NIH award (NICHD-5R24HD000836) from the Eunice Kennedy Shriver National Institute of Child Health and Human Development.

## DATA AVAILABILTY

Sequencing data used in this study is deposited at EMBL-EBI ArrayExpress. Single-cell RNA sequencing of human fetal lung and human fetal bud tip organoids: human fetal lung (ArrayExpress: E-MTAB-8221, ArrayExpress: E-MTAB-10662) (1, 39), CellTagged organoids (this study), airway cultures (this study).

Code used to process data can be found at: https://github.com/jason-spence-lab/Conchola_2022

