## Supplemental Info for "Distinct airway progenitor cells drive epithelial heterogeneity in the developing human lung"

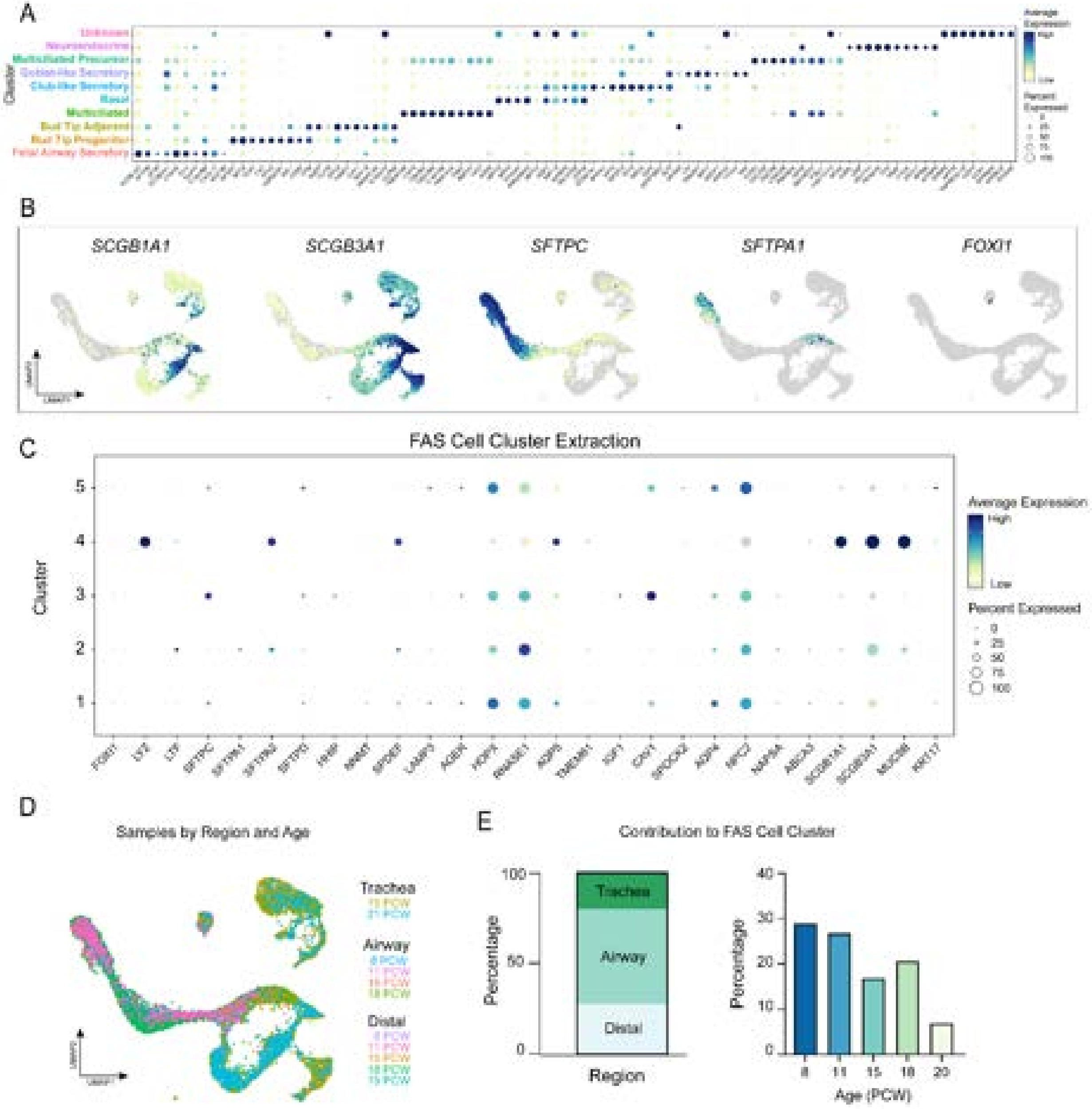

**Supplemental Figure 1: Characterization of FAS cells *in vivo***

(A) Dot plot of top 10 genes defining each cluster from Fig. 1A UMAP cluster plot of fetal lung epithelial data. The dot size represents the percentage of cells expressing the gene in the corresponding cluster, and the dot color indicates log-normalized expression level of the gene. Corresponding clusters are color-coded and labeled along left side of plot.

(B) UMAP feature plot of additional secretory (*SCGB1A1, SCGB3A1*), alveolar (*SFTPC, SFTPA1*), and ionocyte (*FOXI1*) markers. None of these markers are specific to the Fetal Airway Secretory Cluster. The color of each dot in the feature plot indicates log-normalized expression level of the genes in the represented cell.

(C) Dot plot of additional secretory genes from Fig. 1C UMAP cluster plot of Fetal Airway Secretory cell extraction. The dot size represents the percentage of cells expressing the gene in the corresponding cluster, and the dot color indicates log-normalized expression level of the gene.

(D) UMAP cluster plot of fetal lung epithelial data by sample. Each dot represents a single cell and cells were computationally clustered based on transcriptional similarities. 6 biologically distinct samples were sequenced at PCW ages 8, 11, 15, 18, 19, 21 and distinct regions of trachea, airway or distal lung were sequenced among the samples.

(E) Quantification of cells contributing to Fetal Airway Secretory cluster are reported as percentages by region (left panel) or sample age (right panel).

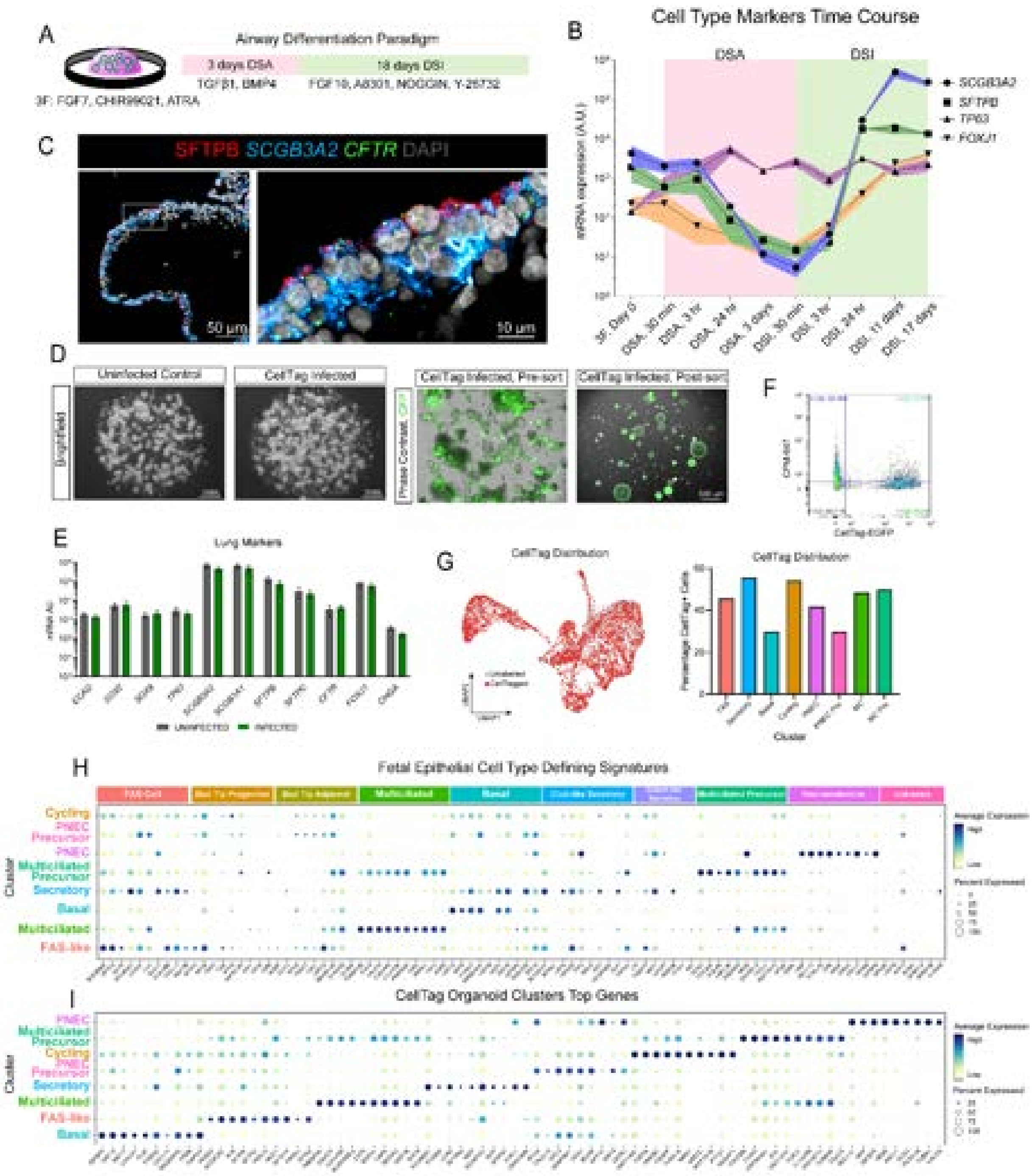

**Supplemental Figure 2: Characterization of 21-day Airway Organoids and CellTagged Cultures**

(A) Overview of Airway Differentiation Paradigm using BTOs: 3F BTOs are treated with 3-day pulse of ‘Dual-SMAD Activation’ (DSA) media containing TGFβ1 and BMP4. On day 4, organoids are treated with ‘Dual-SMAD Inhibition’ (DSI) media containing FGF10, A-8301, NOGGIN, and Y-26732 for 18 more days.

(B) Time course RT-qPCR data of specified cell markers over 21-day Airway Differentiation including basal cell marker *TP63*, FAS cell markers *SCGB3A2* and *SFTPB*, and multiciliated cell marker *FOXJ1*. Bands represent error bars as standard error of mean. This quantification was performed on three technical replicates from three biological distinct samples.

(C) Representative FISH with co-IF image of 21-day Airway Organoid expressing FAS cell markers *SCGB3A2, CFTR*, and SFTPB.

(D) Representative brightfield images of uninfected control and CellTagged-Airway Organoids (top). Representative fluorescence images of CellTagged cultures for GFP expression pre- (7d post-infection) and post-sort (14d post-sort).

(E) RT-qPCR data comparing expression of lung epithelial markers *ECAD, SOX2, SOX9, TP63*, *SCGB3A2, SCGB1A1, SFTPB, SFTPC, CFTR, FOXJ1,* and *CHGA,* between uninfected (grey) and infected (green) Airway Organoids 7 days post-infection. This quantification was performed on three separate biological replicates with three technical replicates each. Error bars represent standard error of the mean. Statistical tests were performed by Welch’s t-test (unpaired, two-tailed). P values > 0.14 for comparison of conditions among all genes.

(F) Representative gating plot for fluorescence activated cell sorting (FACS) of CellTagged-Airway Organoids, 7 days after infection for CellTag-EGFP and CPM-AF647 expression.

(G) UMAP plot of all sequenced cells from CellTagging experiment. Red dots label cells that contained CellTag barcode (left). Histogram of percentage of CellTag-expressing cells from each cluster, labeled and color-coded according to Fig. 2C.

(H) Dot plot of top 10 Fetal Epithelial Cell Type Defining Signatures from Fig. 1A UMAP cluster plot used to define the cell type clusters of 58-day CellTagged-Airway Organoids. Fetal Epithelial Signatures are defined along the top of the plot and cell type clusters for Fig. 2C are labeled accordingly on left side of plot. The dot size represents the percentage of cells expressing the gene in the corresponding cluster, and the dot color indicates log-normalized expression level of the gene. Complete gene lists for cell scoring are in Table S2.

(I) Dot plot of top 10 genes from each cluster of 58-day CellTagged-Airway Organoids. The dot size represents the percentage of cells expressing the gene in the corresponding cluster, and the dot color indicates log-normalized expression level of the gene. Corresponding clusters are color-coded and labeled along left side of plot.

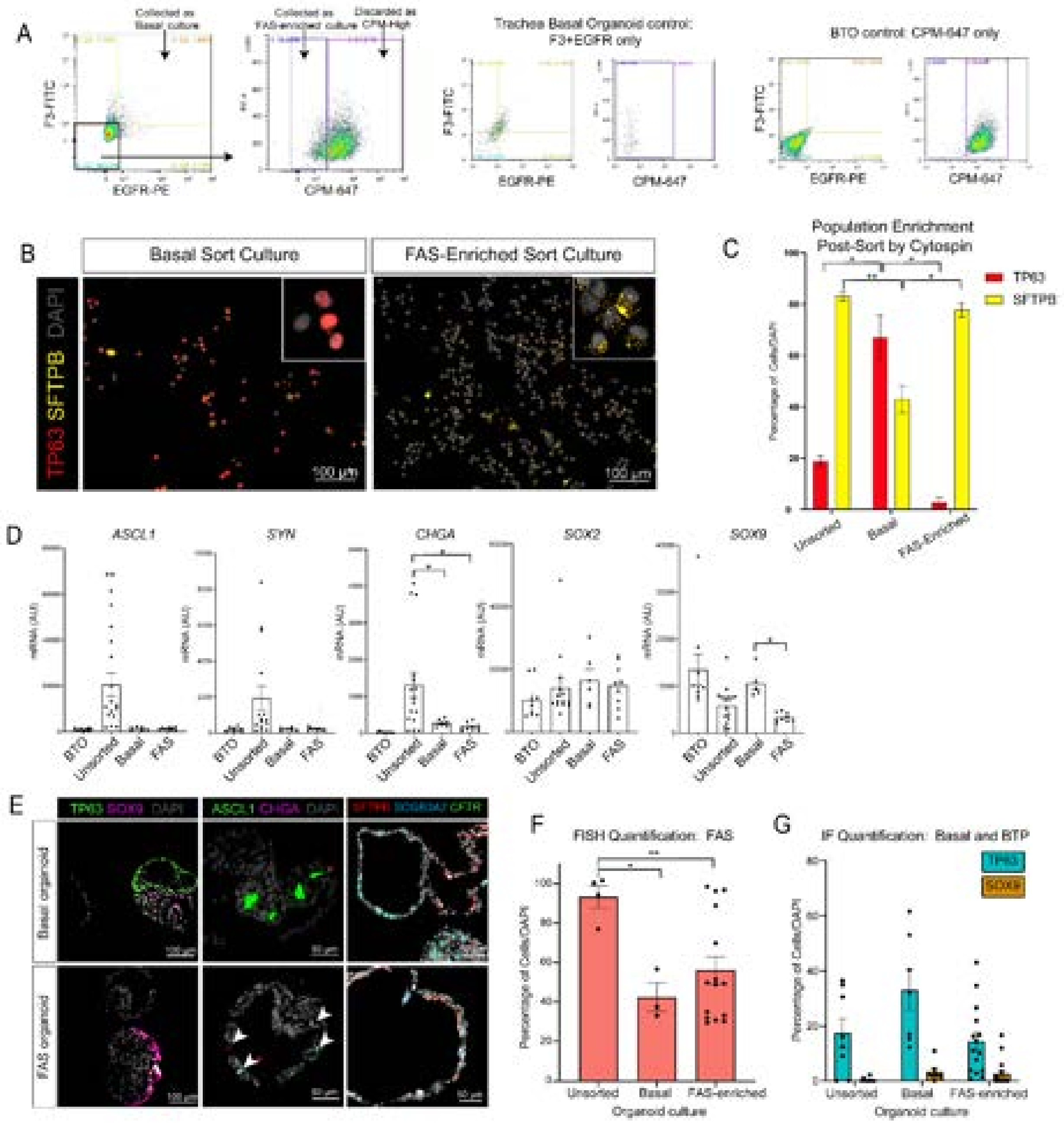

**Supplemental Figure 3: Validation of FAS sorting strategy and enriched organoid cultures**

(A) Representative gating plots for FACS for Fig. 4A. Basal cells were sorted using F3-FITC and/or EGFR-PE conjugated antibodies. Remaining negative culture was then sorted for CPM expression, and high-expressing cells were removed as Bud Tip Progenitors leaving a triplenegative population of 'FAS-enriched' cells. Representative gating controls using F3+EGFR on tracheal basal organoids and CPM on 3F BTOs.

(B) Immunofluorescence images of basal and FAS-enriched cultures processed by Cytospin immediately following cell sorting. Slides were stained for TP63 (basal cell) and SFTPB (FAS cell) to confirm enrichment of the 2 populations after sorting.

(C) Quantification of $TP63^+$ and $SFTPB^+$ cells as percentage of $DAPI^+$ cells on Cytospin-processed cultures. This quantification was performed on three imaged regions of samples across two experiments. Statistical tests were performed by Welch's t-test (unpaired, two-tailed) comparing TP63 to SFTPB for each condition; p-values < 0.0001 (Unsorted), = 0.0625 (Basal), <0.0001 (FAS). For comparison of each marker across conditions, one-way ANOVA with Welch's correction was used; p-values are (*) <0.05, (**) <0.005.

(D) RT-qPCR data comparing expression of PNEC markers (*ASCL1, SYN, CHGA*) including unsorted condition; airway marker *SOX2* and bud tip marker (*SOX9*). This quantification was performed on three technical replicates from two to three biological replicates per condition. Error bars represent standard error of the mean. Statistical tests were performed by one-way ANOVA with Welch's correction; p-values are (*) <0.05, (**) <0.05.

(E) FISH and co-IF staining on paraffin sections of basal and FAS-enriched organoids 2.5 weeks post-sort for basal (TP63), bud tip (SOX9), PNEC (ASCL1, CHGA), and FAS (*SCGB3A2, SFTPB, CFTR*) cell types.

(F) Quantification of $SFTPB^+$/$\textit{SCGB3A2}^+$/$\textit{CFTR}^+$ cells in unsorted, basal, and FAS-enriched organoids from images represented in Fig. S3E. This quantification was performed on three technical replicates from one to three biological replicates per condition. Error bars represent standard error of the mean. Statistical tests were performed by one-way ANOVA with Welch's correction; p-values are (*) <0.05, (**) <0.005, 0.4860 (Basal-FAS).

(G) Quantification of $TP63^+$ and $SOX9^+$ cells in unsorted, basal, and FAS-enriched organoids from images represented in Fig. S3E. This quantification was performed on three technical replicates from one to three biological replicates per condition. Error bars represent standard error of the mean. Statistical tests were performed by one-way ANOVA with Welch's correction; p-values > 0.12 across all conditions for BTPs and Basal cells.

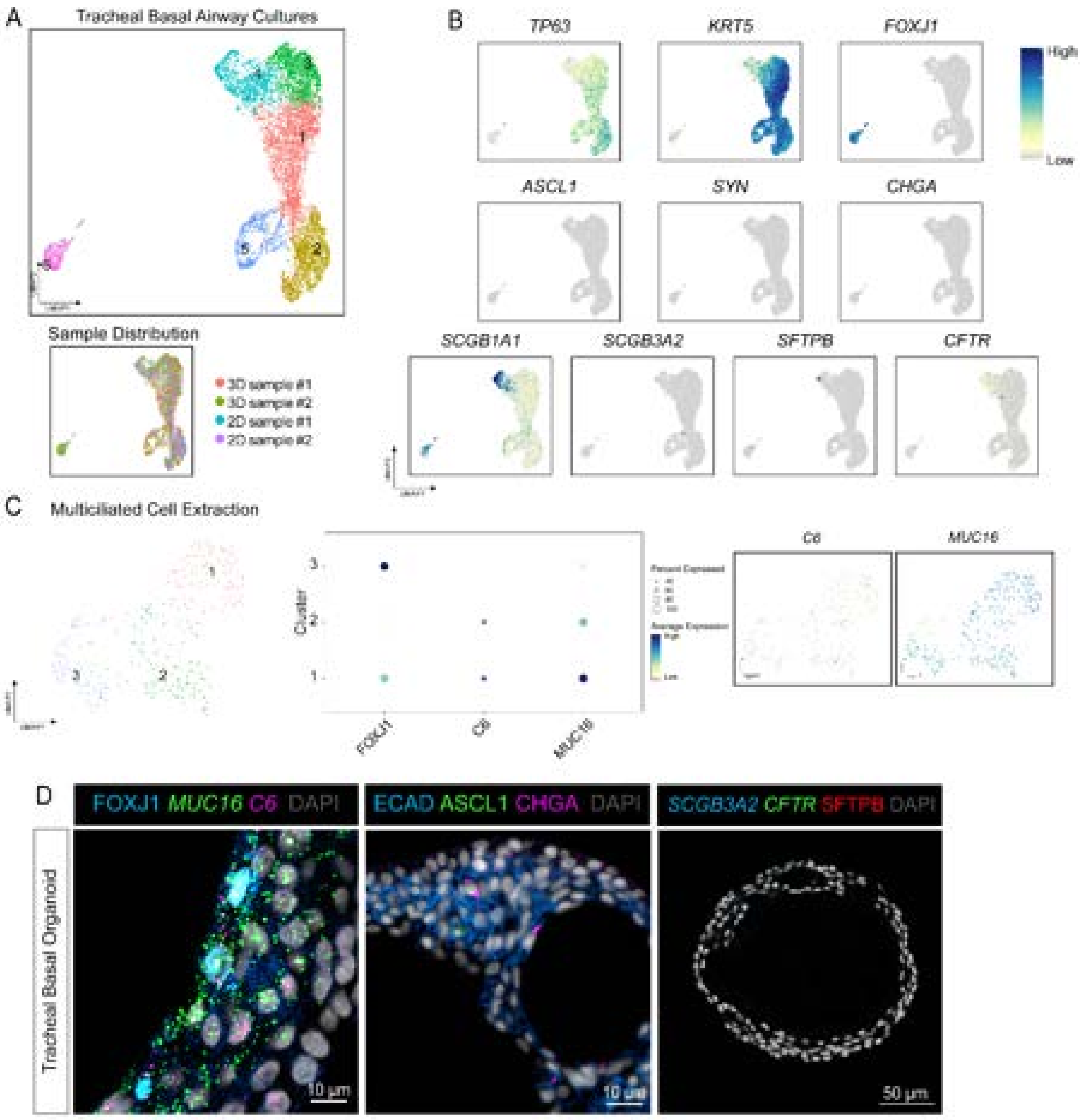

**Supplemental Figure 4: Tracheal basal airway organoids support basal lineages**

(A) UMAP cluster plot of tracheal basal airway. Each dot represents a single cell and cells were computationally clustered based on transcriptional similarities. 2 biologically distinct samples were sequenced from both culture methods: 2D air-liquid interface and 3D organoid cultures.

(B) Feature plots of epithelial cell markers of interest within tracheal basal organoids including basal cell (*TP63, KRT5*), multiciliated (*FOXJ1*), neuroendocrine cell markers (*ASCL1, SYN, CHGA*), and secretory markers (*SCGB1A1, SCGB3A2, SFTPB, CFTR*). The color of each dot in the feature plot indicates log-normalized expression level of the genes in the represented cell.

(C) UMAP cluster plot of multiciliated cell (cluster 6) extraction from tracheal basal airway cultures in Fig. S4A. The multiciliated cell cluster was computationally extracted and re-clustered resulting in 3 subclusters. The plot is colored and numbered by cluster. Middle panel is a dot plot for expression of *FOXJ1*, *C6*, and *MUC16* within multiciliated cell extraction. The dot size represents the percentage of cells expressing the gene in the corresponding cluster and the color indicates log-normalized expression level of the gene. Feature plots for *C6* and *MUC16* on multiciliated cell extraction plot. The color of each dot in the feature plot indicates log-normalized expression level of the genes in the represented cell.

(D) FISH and co-IF staining on paraffin sections of tracheal basal organoids representative for multiciliated (*C6*, *MUC16*, FOXJ1), PNEC (ASCL1, CHGA), and FAS cell (*SCGB3A2, CFTR*, SFTPB) markers.

**Supplemental Table 1: Top 100 FAS Cluster Enriched Genes**

| | p_val | avg_log2F( | pct.1 | pct.2 |
|---|---|---|---|---|
| SCGB3A2 | 0 | 5.254955 | 0.995 | 0.914 |
| SFTPB | 0 | 2.843478 | 0.742 | 0.398 |
| SCGB3A1 | 2.63E-06 | 2.354496 | 0.643 | 0.569 |
| CYB5A | 0 | 1.895231 | 0.982 | 0.851 |
| CLU | 3.50E-262 | 1.805737 | 0.862 | 0.668 |
| VIM | 0 | 1.790674 | 0.899 | 0.603 |
| KLK11 | 4.07E-229 | 1.60825 | 0.5 | 0.207 |
| C16orf89 | 0 | 1.397534 | 0.73 | 0.396 |
| CXCL17 | 0 | 1.363571 | 0.871 | 0.434 |
| MUC5B | 1.36E-107 | 1.323392 | 0.261 | 0.097 |
| XBP1 | 0 | 1.193944 | 0.861 | 0.629 |
| MGST1 | 0 | 1.168161 | 0.945 | 0.77 |
| SSR4 | 0 | 1.153098 | 0.942 | 0.893 |
| CFH | 0 | 1.138915 | 0.638 | 0.25 |
| C3 | 1.41E-287 | 1.132848 | 0.316 | 0.056 |
| CYTL1 | 7.98E-225 | 1.098291 | 0.33 | 0.079 |
| CP | 2.86E-253 | 1.07395 | 0.536 | 0.208 |
| TFF3 | 5.64E-96 | 1.041967 | 0.769 | 0.528 |
| CFHR1 | 3.98E-200 | 1.016482 | 0.275 | 0.058 |
| TMEM45A | 3.17E-134 | 0.968329 | 0.367 | 0.158 |
| RNASE1 | 7.24E-188 | 0.949518 | 0.809 | 0.535 |
| LMO3 | 0 | 0.915306 | 0.456 | 0.109 |
| SERPINF1 | 1.29E-187 | 0.911362 | 0.612 | 0.332 |
| STEAP4 | 2.24E-275 | 0.904796 | 0.395 | 0.102 |
| RNU12 | 1.79E-80 | 0.858121 | 0.519 | 0.34 |
| HNMT | 1.47E-212 | 0.833802 | 0.65 | 0.357 |
| SPINK5 | 1.54E-114 | 0.826323 | 0.301 | 0.117 |
| TMSB4X | 8.51E-166 | 0.777414 | 0.999 | 0.998 |
| FMO2 | 1.28E-255 | 0.769676 | 0.299 | 0.055 |
| COL1A1 | 1.81E-29 | 0.759403 | 0.509 | 0.416 |
| PTP4A1 | 1.90E-109 | 0.736011 | 0.661 | 0.504 |
| SERPINA1 | 1.93E-132 | 0.716498 | 0.416 | 0.185 |
| NDRG1 | 1.27E-155 | 0.714321 | 0.651 | 0.372 |
| PTP4A2 | 3.99E-128 | 0.680505 | 0.629 | 0.428 |
| SFTA2 | 3.34E-97 | 0.669593 | 0.442 | 0.237 |
| SLC4A4 | 1.03E-195 | 0.668346 | 0.352 | 0.107 |
| KLHL24 | 1.00E-153 | 0.667901 | 0.492 | 0.246 |
| IFITM3 | 2.78E-98 | 0.662632 | 0.813 | 0.707 |
| SERPINI1 | 2.54E-116 | 0.653671 | 0.255 | 0.083 |
| CFTR | 1.46E-236 | 0.652506 | 0.324 | 0.072 |
| GDF15 | 8.25E-142 | 0.645965 | 0.532 | 0.25 |
| IFITM2 | 8.18E-97 | 0.64492 | 0.546 | 0.35 |
| SOD3 | 6.25E-111 | 0.644158 | 0.431 | 0.217 |
| SLC39A6 | 3.75E-77 | 0.612614 | 0.474 | 0.31 |
| RAMP2 | 1.32E-89 | 0.611966 | 0.42 | 0.242 |

| | | | | |
|---|---|---|---|---|
| COL1A2 | 4.97E-31 | 0.590732 | 0.473 | 0.36 |
| HOPX | 2.32E-195 | 0.580222 | 0.643 | 0.344 |
| COL3A1 | 1.03E-23 | 0.565608 | 0.38 | 0.282 |
| MGP | 4.40E-41 | 0.555186 | 0.311 | 0.185 |
| ASAH1 | 6.59E-87 | 0.526774 | 0.846 | 0.768 |
| KIAA1324 | 1.21E-141 | 0.50867 | 0.386 | 0.151 |
| LGALS3BP | 5.14E-54 | 0.478135 | 0.7 | 0.625 |
| MET | 3.16E-128 | 0.467998 | 0.318 | 0.119 |
| DHCR24 | 2.67E-68 | 0.460805 | 0.466 | 0.293 |
| AGR2 | 1.25E-75 | 0.454465 | 0.834 | 0.665 |
| NDUFA4L2 | 2.17E-64 | 0.451757 | 0.253 | 0.116 |
| MDK | 2.16E-65 | 0.44767 | 0.977 | 0.888 |
| EFCAB4A | 1.61E-69 | 0.434754 | 0.539 | 0.355 |
| DSTN | 1.32E-80 | 0.434227 | 0.943 | 0.942 |
| CST3 | 2.08E-139 | 0.428529 | 0.961 | 0.921 |
| HEXIM1 | 3.66E-33 | 0.419454 | 0.445 | 0.345 |
| CRLF1 | 6.44E-26 | 0.414759 | 0.29 | 0.202 |
| PPIB | 8.30E-59 | 0.408622 | 0.881 | 0.842 |
| HMGB3 | 6.21E-12 | 0.402339 | 0.431 | 0.393 |
| FAM46C | 1.24E-75 | 0.401334 | 0.29 | 0.139 |
| CD55 | 4.00E-36 | 0.393025 | 0.505 | 0.394 |
| SPARCL1 | 6.07E-17 | 0.38617 | 0.346 | 0.275 |
| HLA-A | 4.52E-45 | 0.384311 | 0.694 | 0.603 |
| BMP3 | 2.05E-17 | 0.383772 | 0.409 | 0.346 |
| CCDC80 | 6.54E-10 | 0.382432 | 0.264 | 0.211 |
| NPNT | 7.78E-75 | 0.380305 | 0.367 | 0.196 |
| TSPAN4 | 7.94E-45 | 0.37018 | 0.458 | 0.324 |
| HES4 | 1.25E-53 | 0.369087 | 0.541 | 0.365 |
| WSB1 | 4.12E-26 | 0.368924 | 0.796 | 0.771 |
| TMEM150( | 6.31E-50 | 0.366687 | 0.296 | 0.168 |
| HSPA5 | 7.97E-33 | 0.360747 | 0.765 | 0.734 |
| GRN | 3.95E-36 | 0.354947 | 0.611 | 0.529 |
| SUSD4 | 5.15E-40 | 0.329044 | 0.303 | 0.19 |
| PPIC | 1.37E-24 | 0.318208 | 0.475 | 0.388 |
| TMEM205 | 1.86E-30 | 0.317163 | 0.652 | 0.582 |
| CFI | 1.54E-45 | 0.317028 | 0.323 | 0.19 |
| PLK2 | 5.57E-16 | 0.31405 | 0.337 | 0.266 |
| LY6E | 5.21E-30 | 0.313576 | 0.74 | 0.675 |
| SOCS3 | 2.58E-28 | 0.309885 | 0.527 | 0.412 |
| EMID1 | 1.58E-28 | 0.307437 | 0.327 | 0.234 |
| CHN1 | 5.45E-47 | 0.304749 | 0.25 | 0.133 |
| NAMPT | 2.43E-23 | 0.296069 | 0.292 | 0.209 |
| RHOB | 1.85E-16 | 0.294159 | 0.657 | 0.618 |
| ID4 | 5.20E-18 | 0.278054 | 0.382 | 0.309 |
| GADD45B | 4.26E-18 | 0.271855 | 0.723 | 0.656 |
| GRHL1 | 4.03E-35 | 0.266027 | 0.299 | 0.189 |
| NEDD4L | 4.34E-15 | 0.261845 | 0.378 | 0.317 |

| | | | | |
|---|---|---|---|---|
| LINC00910 | 9.68E-09 | 0.258019 | 0.259 | 0.211 |
| FGFR3 | 8.20E-17 | 0.25733 | 0.372 | 0.295 |
| NENF | 6.71E-16 | 0.252012 | 0.627 | 0.567 |
| FKBP2 | 1.56E-22 | 0.251541 | 0.775 | 0.752 |
| PRDX4 | 5.71E-15 | 0.251453 | 0.622 | 0.586 |
| LTA4H | 2.50E-15 | 0.251124 | 0.536 | 0.485 |
| PLD3 | 4.31E-29 | 0.250696 | 0.829 | 0.793 |
| NR2F2 | 3.89E-34 | -0.25004 | 0.226 | 0.359 |

**Supplemental Table 2: Cell Scoring Gene Lists**

| **Secretory** | **Multiciliat** | **PNEC** | **FAS** | **MC Precur** | **Basal** |
|---|---|---|---|---|---|
| SCGB1A1 | TMEM190 | GHRL | SCGB3A2 | CDC20B | KRT15 |
| KRT4 | CAPS | GRP | SFTPB | LRRC26 | KRT5 |
| WFDC2 | C9orf24 | SEC11C | CYB5A | HES6 | S100A2 |
| MSLN | C20orf85 | PCSK1N | VIM | ZMYND10 | MIR205HG |
| IGF2 | C1orf194 | CPE | C16orf89 | CDK1 | AQP3 |
| SERPINB3 | FAM183A | NNAT | CXCL17 | CCDC74A | KRT19 |
| SLPI | RP11-356K | CHGA | XBP1 | CCDC67 | TACSTD2 |
| ANXA1 | OMG | ASCL1 | MGST1 | HYLS1 | SFN |
| MUC4 | RSPH1 | RPRM | SSR4 | NEK2 | JUNB |
| S100A9 | PIFO | IGFBP5 | CFH | CDC20 | KLF5 |
| KRT13 | AGR3 | HES6 | LMO3 | C1orf192 | PERP |
| S100P | ODF3B | BEX1 | C3 | PLK4 | F3 |
| KRT19 | SNTN | ACSL1 | STEAP4 | SPAG6 | CD9 |
| TACSTD2 | FABP6 | SCGN | CLU | RIBC2 | HCAR3 |
| UPK1B | TSPAN1 | HEPACAM2 | FMO2 | CCDC19 | IGFBP2 |
| IGFBP3 | CCDC78 | CALCA | CP | ANLN | JUN |
| S100A6 | DYNLRB2 | MEG3 | CFTR | POC1A | CSTA |
| AQP3 | DNAAF1 | TMEM176I | KLK11 | HIST1H2BJ | NPPC |
| CLDN4 | TPPP3 | TMEM176/ | CYTL1 | FOXN4 | HCAR2 |
| APOBEC3A | C5orf49 | BIK | HNMT | E2F7 | FOS |
| CYP2F1 | MS4A8 | SCG5 | CFHR1 | MCIDAS | IL33 |
| RHOV | C9orf116 | APOA1BP | SLC4A4 | SYT5 | DLK2 |
| TNNT3 | CAPSL | UCHL1 | HOPX | STIL | ADH7 |
| CEACAM6 | SMIM22 | MIAT | RNASE1 | C11orf88 | CAPNS2 |
| C19orf33 | C9orf117 | C12orf75 | SERPINF1 | CCNO | PKP1 |
| PLAC8 | CETN2 | KLK12 | TMSB4X | C5orf49 | TP63 |
| KLF5 | AC013264. | CHGB | NDRG1 | MELK | PNCK |
| ELF3 | MORN5 | SLC35D3 | KLHL24 | ROPN1L | RPLP1 |
| FAM3D | TSPAN19 | NPDC1 | GDF15 | RP11-263K | BHLHE40 |
| LSP1 | TUBB4B | DDC | KIAA1324 | C11orf16 | HSPB1 |
| VMO1 | CRIP1 | TAGLN3 | CST3 | SGOL2 | ZFP36 |
| F3 | CDHR3 | GFRA3 | TMEM45A | MUC12 | GJB2 |
| LCN2 | MRPS31 | SCG3 | SERPINA1 | FOXJ1 | KRT13 |
| LYPD2 | MORN2 | NPW | MET | FAM183A | LSP1 |
| CSTA | C1orf192 | SCG2 | PTP4A2 | C7orf57 | EGR1 |
| ASS1 | FAM92B | TUBB2B | SERPINI1 | TEKT2 | IGFBP3 |
| SLC16A9 | PSENEN | C3orf14 | SPINK5 | DNAH12 | SERPINB5 |
| TNFSF10 | FAM216B | ATP6V0B | SOD3 | RSPH9 | IER3 |
| SERPINB2 | TMC5 | MAOB | PTP4A1 | STMND1 | ARL4D |
| GPR110 | ZMYND10 | VAMP2 | MUC5B | LRRC46 | MYC |
| MUC20 | CCDC17 | C4orf48 | IFITM3 | DNAAF3 | MPZL2 |
| EPHA2 | LRRIQ1 | APLP1 | SFTA2 | PITPNM1 | COL7A1 |
| TCN1 | C22orf15 | LINC00261 | IFITM2 | KDELC2 | PLP2 |
| A4GALT | C21orf58 | BEX2 | TFF3 | RSPH1 | MEG3 |
| GABRP | C11orf88 | BCAM | RAMP2 | ARMC3 | ZFP36L1 |

| | | | | | |
|---|---|---|---|---|---|
| LYN | EFHC1 | LY6H | ASAH1 | CEP152 | CALML3 |
| PDE4C | SLC44A4 | CA8 | DSTN | MYB | FOSB |
| FUT3 | LRRC46 | FAM105A | RNU12 | PIFO | LMNA |
| TFF3 | CES1 | TUBA1A | SLC39A6 | SPEF1 | PVRL1 |
| FABP5 | FOXJ1 | HOXB5 | FAM46C | CAPSL | ACKR3 |

**Supplemental Table 3: Differential Gene Expression: CellTag Multiciliated Extraction**

| **Cluster 1** | p_val | avg_log2FC | pct.1 | pct.2 |
|---|---|---|---|---|
| TMSB4X | 5.44E-72 | 1.913272 | 1 | 0.989 |
| RPS12 | 1.19E-59 | 1.887726 | 1 | 0.989 |
| ZFAS1 | 2.74E-66 | 1.646498 | 0.996 | 0.947 |
| TMSB10 | 2.02E-70 | 1.560005 | 1 | 0.979 |
| ACTG1 | 2.34E-60 | 1.5524 | 1 | 0.963 |
| RPS3 | 1.62E-67 | 1.541739 | 1 | 0.981 |
| RPL10 | 1.46E-60 | 1.506078 | 1 | 0.997 |
| RPS18 | 1.22E-63 | 1.452995 | 1 | 1 |
| RPL36A | 3.07E-61 | 1.446298 | 1 | 0.971 |
| RPL12 | 8.25E-67 | 1.361668 | 1 | 0.992 |
| TPT1 | 1.26E-74 | 1.356313 | 1 | 0.987 |
| RPS2 | 1.68E-74 | 1.348675 | 1 | 0.984 |
| GAS5 | 1.87E-62 | 1.338472 | 0.92 | 0.54 |
| RPS5 | 1.14E-63 | 1.317728 | 1 | 0.971 |
| NDRG1 | 1.76E-60 | 1.30898 | 0.837 | 0.238 |
| RPLP0 | 5.09E-76 | 1.258293 | 1 | 0.966 |
| RPL5 | 2.85E-70 | 1.21394 | 1 | 0.981 |
| RPL14 | 1.01E-68 | 1.212795 | 1 | 0.979 |
| RPL32 | 2.95E-63 | 1.206642 | 1 | 0.987 |
| NACA | 5.15E-68 | 1.199075 | 1 | 0.984 |
| RPS23 | 9.99E-59 | 1.195777 | 1 | 0.989 |
| RPL18 | 7.83E-69 | 1.186185 | 1 | 0.989 |
| RPL26 | 5.32E-67 | 1.151449 | 1 | 0.989 |
| PABPC1 | 4.80E-65 | 1.122136 | 0.996 | 0.907 |
| RACK1 | 2.41E-61 | 1.115608 | 1 | 0.989 |
| EEF2 | 1.24E-66 | 1.093645 | 0.996 | 0.921 |
| RPS6 | 4.69E-63 | 1.083167 | 1 | 0.984 |
| RPS19 | 3.00E-59 | 1.068597 | 1 | 0.997 |
| RPS3A | 1.92E-62 | 1.065055 | 1 | 0.992 |
| RPS21 | 1.64E-54 | 1.022221 | 1 | 0.987 |
| RPL13A | 1.61E-66 | 1.010743 | 0.996 | 0.974 |
| PPP1R14B | 7.16E-57 | 0.999735 | 0.943 | 0.614 |
| RPL28 | 1.93E-59 | 0.985805 | 1 | 0.987 |
| RPL35 | 2.47E-58 | 0.976773 | 1 | 0.981 |
| NPM1 | 1.52E-60 | 0.973048 | 1 | 0.971 |
| RPL24 | 1.49E-63 | 0.96525 | 1 | 0.989 |
| RPL7A | 3.27E-59 | 0.963441 | 1 | 0.997 |
| RPL37 | 1.12E-60 | 0.953439 | 1 | 0.992 |
| RPL18A | 6.01E-61 | 0.944108 | 1 | 0.997 |
| RPL34 | 8.83E-63 | 0.942735 | 1 | 1 |
| RPL30 | 8.29E-61 | 0.937351 | 1 | 0.987 |
| RPL13 | 5.14E-61 | 0.937189 | 1 | 1 |
| RPL6 | 3.42E-58 | 0.934455 | 1 | 0.984 |
| RPL8 | 2.89E-54 | 0.927118 | 1 | 0.984 |
| MARCKSL1 | 1.77E-61 | 0.923198 | 0.826 | 0.241 |

| **Cluster 2** | p_val | avg_log2FC |
|---|---|---|
| ALOX15 | 1.72E-41 | 1.501288 |
| CD24 | 9.29E-49 | 1.230938 |
| C20orf85 | 2.31E-47 | 1.078671 |
| AC013264. | 2.99E-44 | 1.032638 |
| MUC16 | 8.05E-38 | 1.030274 |
| BAALC | 1.19E-40 | 0.998926 |
| TUBA1A | 7.70E-35 | 0.994367 |
| CAPSL | 2.86E-46 | 0.985319 |
| PLAAT2 | 5.86E-38 | 0.982311 |
| SLAIN2 | 1.75E-42 | 0.935534 |
| WWOX-AS: | 4.13E-33 | 0.905017 |
| DYNLRB2 | 1.60E-51 | 0.898855 |
| CALM1 | 2.05E-44 | 0.893163 |
| SPA17 | 3.72E-46 | 0.880303 |
| ROPN1L | 6.20E-45 | 0.879038 |
| GSTP1 | 3.68E-53 | 0.870336 |
| MORN5 | 4.63E-42 | 0.867185 |
| AL357093.: | 9.77E-34 | 0.864698 |
| C9orf24 | 1.84E-42 | 0.861729 |
| WDR38 | 1.13E-42 | 0.841744 |
| ANKUB1 | 1.72E-38 | 0.830752 |
| C1orf194 | 1.80E-42 | 0.828788 |
| PRDX5 | 4.21E-47 | 0.820879 |
| SNTN | 2.55E-37 | 0.820119 |
| NME5 | 6.36E-37 | 0.819656 |
| BX005040. | 2.95E-33 | 0.811777 |
| PIFO | 1.35E-52 | 0.807779 |
| TSPAN19 | 1.24E-37 | 0.805986 |
| CFAP300 | 2.93E-36 | 0.805273 |
| MORN2 | 8.08E-43 | 0.793878 |
| CABCOCO1 | 1.03E-39 | 0.789471 |
| PIH1D3 | 2.73E-41 | 0.784606 |
| DNPH1 | 4.95E-39 | 0.778413 |
| UFC1 | 1.16E-44 | 0.77054 |
| TCTEX1D1 | 2.35E-34 | 0.766688 |
| FAM229B | 3.47E-43 | 0.762537 |
| ANKRD66 | 7.47E-35 | 0.759269 |
| DYNLL1 | 1.99E-45 | 0.758259 |
| PKIG | 6.56E-36 | 0.757141 |
| ATP5IF1 | 3.33E-38 | 0.748012 |
| CAPS | 6.06E-44 | 0.734851 |
| RSPH1 | 1.83E-36 | 0.730272 |
| CCDC78 | 3.34E-36 | 0.729948 |
| C11orf74 | 9.34E-38 | 0.72522 |
| FAM92B | 2.55E-34 | 0.709496 |

| | | | | | | | |
|---|---|---|---|---|---|---|---|
| RPL29 | 1.57E-56 | 0.921525 | 1 | 0.997 | ODF3B | 2.78E-35 | 0.707933 |
| RPS15A | 9.43E-60 | 0.898188 | 1 | 0.995 | SMIM41 | 3.10E-44 | 0.695528 |
| EEF1D | 1.71E-54 | 0.89762 | 0.996 | 0.929 | CFAP52 | 1.51E-33 | 0.693084 |
| RPS14 | 6.39E-66 | 0.873561 | 1 | 0.997 | EFCAB1 | 4.79E-36 | 0.674513 |
| RPL39 | 4.58E-56 | 0.858793 | 1 | 0.997 | DNALI1 | 2.83E-39 | 0.674456 |
| EEF1A1 | 2.08E-61 | 0.852702 | 1 | 1 | CIB1 | 2.50E-33 | 0.66112 |
| RPL7 | 7.98E-55 | 0.84586 | 0.996 | 0.958 | ARL3 | 2.18E-34 | 0.657843 |
| RPL15 | 5.23E-63 | 0.823096 | 1 | 0.995 | POLR2I | 1.97E-34 | 0.644039 |
| RPL23A | 2.82E-60 | 0.81936 | 1 | 0.981 | TMEM14B | 8.86E-36 | 0.598291 |
| RPL17 | 3.65E-60 | 0.80588 | 1 | 0.989 | FGF14 | 2.83E-43 | 0.596616 |
| RPL4 | 1.68E-54 | 0.764435 | 0.996 | 0.976 | CETN2 | 9.13E-34 | 0.593724 |
| RPL22 | 2.52E-55 | 0.758492 | 1 | 0.987 | FAM81B | 1.50E-32 | 0.577076 |
| RPS16 | 1.07E-57 | 0.75241 | 1 | 0.981 | SOD1 | 6.34E-36 | 0.576061 |
| RPL9 | 6.22E-59 | 0.749131 | 1 | 0.992 | TSTD1 | 1.15E-34 | 0.572195 |
| RPS27A | 5.60E-55 | 0.69973 | 1 | 0.997 | SEM1 | 2.62E-33 | 0.536849 |

| pct.1 | pct.2 |
|---|---|
| 0.829 | 0.401 |
| 0.973 | 0.977 |
| 0.977 | 0.932 |
| 0.981 | 0.88 |
| 0.822 | 0.432 |
| 0.818 | 0.401 |
| 1 | 0.99 |
| 0.977 | 0.932 |
| 0.795 | 0.385 |
| 0.965 | 0.88 |
| 0.744 | 0.362 |
| 0.992 | 0.982 |
| 1 | 1 |
| 0.984 | 0.977 |
| 0.992 | 0.971 |
| 1 | 1 |
| 0.984 | 0.956 |
| 1 | 0.961 |
| 0.996 | 0.997 |
| 0.953 | 0.943 |
| 0.771 | 0.339 |
| 0.992 | 0.99 |
| 1 | 1 |
| 0.996 | 0.958 |
| 0.961 | 0.885 |
| 0.857 | 0.612 |
| 1 | 1 |
| 0.713 | 0.276 |
| 0.981 | 0.958 |
| 1 | 0.997 |
| 0.969 | 0.901 |
| 0.938 | 0.846 |
| 0.984 | 0.974 |
| 0.988 | 0.997 |
| 0.919 | 0.823 |
| 0.996 | 0.995 |
| 0.938 | 0.883 |
| 1 | 1 |
| 0.915 | 0.766 |
| 0.984 | 1 |
| 1 | 1 |
| 0.996 | 0.997 |
| 0.984 | 0.971 |
| 0.953 | 0.935 |
| 0.992 | 0.964 |

| **Cluster 3** | p_val | avg_log2FC | pct.1 | pct.2 |
|---|---|---|---|---|
| AGR2 | 6.64E-30 | 1.516259 | 0.992 | 0.956 |
| EVL | 3.17E-26 | 1.293999 | 0.958 | 0.768 |
| SMIM6 | 1.07E-26 | 1.244827 | 0.958 | 0.762 |
| MS4A8 | 1.41E-32 | 1.118316 | 0.983 | 0.808 |
| FOLR1 | 5.79E-27 | 1.056022 | 0.875 | 0.567 |
| AGR3 | 2.92E-38 | 0.973058 | 1 | 0.977 |
| C1orf189 | 1.24E-23 | 0.956813 | 1 | 0.831 |
| UGT2B17 | 1.82E-37 | 0.941825 | 0.725 | 0.195 |
| CDHR3 | 5.00E-23 | 0.931962 | 0.992 | 0.828 |
| RSPH9 | 3.20E-31 | 0.910698 | 1 | 0.975 |
| C22orf15 | 2.08E-26 | 0.904455 | 1 | 0.92 |
| KIF21A | 4.63E-28 | 0.88738 | 1 | 0.822 |
| C6 | 6.95E-41 | 0.843242 | 0.85 | 0.266 |
| TPPP3 | 1.59E-28 | 0.830736 | 1 | 0.992 |
| SAXO2 | 3.03E-31 | 0.824052 | 1 | 0.91 |
| DYNC2H1 | 1.33E-27 | 0.821865 | 0.992 | 0.879 |
| CATSPERE | 1.43E-28 | 0.780021 | 0.825 | 0.395 |
| STK33 | 4.46E-26 | 0.777669 | 0.992 | 0.879 |
| MLLT1 | 1.34E-23 | 0.763555 | 0.95 | 0.67 |
| C12orf75 | 7.63E-25 | 0.76116 | 1 | 0.946 |
| TCTEX1D4 | 4.95E-20 | 0.729802 | 1 | 0.927 |
| SPACA9 | 1.08E-23 | 0.729073 | 1 | 0.939 |
| CD164L2 | 1.03E-20 | 0.720965 | 1 | 0.92 |
| DTHD1 | 1.50E-19 | 0.720616 | 0.992 | 0.816 |
| CCDC181 | 9.02E-28 | 0.717249 | 0.958 | 0.669 |
| CCDC81 | 1.04E-28 | 0.71546 | 0.917 | 0.504 |
| LDLRAD1 | 5.58E-23 | 0.714515 | 1 | 0.918 |
| C5orf15 | 1.00E-23 | 0.697451 | 1 | 0.92 |
| ZDHHC1 | 4.06E-21 | 0.697359 | 1 | 0.856 |
| C11orf97 | 3.30E-24 | 0.696157 | 1 | 0.964 |
| LINC02345 | 5.31E-26 | 0.680836 | 0.917 | 0.554 |
| GIHCG | 1.47E-20 | 0.673259 | 1 | 0.944 |
| C21orf58 | 1.81E-26 | 0.671654 | 1 | 0.933 |
| EFHC1 | 1.18E-25 | 0.666664 | 1 | 0.946 |
| TMEM107 | 2.18E-24 | 0.663412 | 1 | 0.943 |
| SPAG17 | 8.73E-23 | 0.655861 | 0.992 | 0.887 |
| BSCL2 | 7.28E-23 | 0.65571 | 1 | 0.971 |
| ETFB | 1.15E-19 | 0.651536 | 0.983 | 0.933 |
| GALM | 5.14E-23 | 0.647753 | 0.858 | 0.569 |
| FOXJ1 | 3.40E-20 | 0.644868 | 1 | 0.954 |
| DRC3 | 4.04E-20 | 0.637964 | 0.992 | 0.837 |
| PTPRN2 | 3.62E-19 | 0.63729 | 0.85 | 0.59 |
| IQCD | 4.13E-23 | 0.632368 | 1 | 0.902 |
| KIAA0408 | 7.26E-27 | 0.631008 | 0.8 | 0.366 |
| IFT57 | 2.20E-22 | 0.626875 | 1 | 0.992 |

| | | | | | | |
|---|---|---|---|---|---|---|
| 0.996 | 0.99 | ASL | 4.09E-24 | 0.610365 | 0.933 | 0.824 |
| 0.81 | 0.344 | PALMD | 1.43E-20 | 0.609915 | 0.942 | 0.766 |
| 0.953 | 0.932 | SNTN | 1.57E-19 | 0.609092 | 1 | 0.967 |
| 0.984 | 0.977 | GPC5-AS1 | 3.08E-22 | 0.602779 | 0.867 | 0.552 |
| 0.984 | 0.995 | FAM183A | 5.74E-23 | 0.594663 | 1 | 0.99 |
| 0.992 | 1 | AC130456. | 3.81E-21 | 0.578764 | 0.925 | 0.697 |
| 0.981 | 0.99 | RSPH1 | 2.25E-21 | 0.57741 | 1 | 0.996 |
| 0.992 | 0.997 | TTC26 | 5.29E-20 | 0.560514 | 1 | 0.881 |
| 0.957 | 0.987 | LRRC71 | 1.45E-20 | 0.549624 | 0.975 | 0.816 |
| 0.609 | 0.122 | CFAP73 | 5.93E-20 | 0.548693 | 1 | 0.944 |
| 0.996 | 1 | C17orf97 | 1.72E-19 | 0.545799 | 0.983 | 0.9 |
| 0.961 | 0.977 | AC004832. | 2.62E-19 | 0.532021 | 0.975 | 0.764 |
| 0.996 | 1 | GET1 | 2.23E-19 | 0.516002 | 1 | 0.925 |
| 0.977 | 1 | CST3 | 3.13E-19 | 0.513645 | 1 | 0.964 |
| 0.981 | 0.995 | C6orf141 | 2.67E-22 | 0.470093 | 0.65 | 0.276 |

**Supplementary Table 4. Antibody and TSA Dilutions and Primer Sequences**

| Antibody | Dilution |
|---|---|
| SOX9 (Millipore, Cat#AB5535) | 1:500 |
| CPM (Wako/FujiFilm, Cat#014-27501) | 1:500 for IF; 1:300 for FACS |
| ECAD (R&D, Cat#AF748) | 1:500 |
| ECAD (BD Biosciences, Cat#610181) | 1:500 |
| TP63 (R&D Systems, Cat#BAF1916) | 1:500 |
| CHGA (Santa Cruz Biotechnology, Cat#sc-1488) | 1:100 |
| FOXJ1 (Seven Hills Bioreagents, Cat#WMAB-319) | 1:250 |
| ASCL1 (Santa Cruz Biotechnology, Cat#sc-374104) | 1:50 |
| SP-B (Seven Hills Bioreagents, Cat#WMAB-1B9) | 1:500 |
| SCGB3A2 (Abcam, Cat#ab181853) | 1:50 |
| CD142-FITC, REAfinity (Miltenyi Biotec, Cat#130-115-683) | 1:50 |
| EGFR-PE, REAfinity (Miltenyi Biotec, Cat#130-110-528) | 1:50 |
| **Probe** | **TSA dilution** |
| Hs-SCGB3A2-C1/C2 (ACDbio RNAscope, Cat#549961; Cat#549961-C2) | 1:5000 |
| Hs-SFTPB-C1 (ACDbio RNAscope, Cat#544251) | 1:2500 |
| Hs-CFTR-C1/C2/C3 (ACDbio RNAscope, Cat#603291; Cat#603291-C2; Cat#603291-C3) | 1:2500 |
| Hs-MUC16-C2 (ACDbio RNAscope, Cat#40591-C2) | 1:4000 |

| | |
|---|---|
| Hs-C6 (ACDbio, RNAscope, Cat#429951) | 1:3000 |
| **Primer** | **Sequence** |
| ECAD | F: TTGACGCCGAGAGCTACAC<br>R: GACCGGTGCAATCTTCAAA |
| SOX2 | F: GCTTAGCCTCGTCGATGAAC<br>R: AACCCCAAGATGCACAACTC |
| SOX9 | F: GTACCCGCACTTGCACAAC<br>R:GTGGTCCTTCTTGTGCTGC |
| TP63 | F: CCACAGTACACGAACCTGGG<br>R: CCGTTCTGAATCTGCTGGTCC |
| SFTPC | F: AGCAAAGAGGTCCTGATGGA<br>R: CGATAAGAAGGCGTTTCAGG |
| SFTPB | F: GGGTGTGTGGGACCATGT<br>R: CAGCACTTTAAAGGACGGTGT |
| SCGB3A2 | F: AAGCTGGTAACTATCTTCCTGCT<br>R: AGGGGCACTTTGTTGATGAGG |
| SCGB1A1 | F: ATGAAACTCGCTGTCACCCT<br>R: GTTTCGATGACACGCTGAAA |
| CFTR | F: CCTATGACCCGGATAACAAGGA<br>R: GAACACGGCTTGACAGCTTTA |
| FOXJ1 | F: CAACTTCTGCTACTTCCGCC<br>R: CGAGGCACTTTGATGAAGC |
| C6 | F: CCTGTTTGGGCATGTCTTATTC<br>R: GGGCCTCAGAAGTCACATTAT |
| MUC16 | F: CACCTTCTCCTGTTTCCTCTAC<br>R: CCCATCCTGTCTGTGGTTATC |
| ASCL1 | F: CGAATGGACTTTGGAAGCAG<br>R: CTTTTCTCCCCCTCCCAAC |
| SYN | F: CGAGGTCGAGTTCGAGTACC<br>R: TTCGGCTGACGAGGAGTAGT |
| CHGA | F: CTGTCCTGGCTCTTCTGCTC<br>R: TGACCTCAACGATGCATTTC |
